# A lymphatic-stem cell interactome regulates intestinal stem cell activity

**DOI:** 10.1101/2022.01.29.478341

**Authors:** Rachel E. Niec, Tinyi Chu, Shiri Gur-Cohen, Marina Schernthanner, Lynette Hidalgo, Hilda Amalia Pasolli, Raghu P. Kataru, Babak J. Mehrara, Dana Pe’er, Elaine Fuchs

## Abstract

Barrier epithelia depend on resident stem cells for homeostasis, defense and repair. Intestinal stem cells (ISCs) of the small and large intestines respond to their local microenvironments (niches) to fulfill a continuous demand for tissue turnover, yet the complexity of their niches is still unfolding. Here, we report an extensive lymphatic network that intimately associates with ISCs within these niches. Devising a lymphatic:organoid coculture system, we show that lymphatic-secreted factors maintain ISCs while inhibiting precocious differentiation. Employing a new deconvolution algorithm, BayesPrism, to pair single-cell and spatial transcriptomics, we cartograph the lymphatic ligand:ISC receptor interactomes at high resolution. We unearth crypt lymphatics as a major source of WNT-signaling factors (WNT2, R-SPONDIN-3) known to drive ISC behavior, and REELIN, a hitherto unappreciated ISC regulator secreted by crypt lymphatics. Together, our studies expose lymphatics as a central hub for niche factors that govern the regenerative potential of ISCs.

## Introduction

At the interface between our body and the external environment, barrier epithelial cells must be rejuvenated constantly (Gehart and Clevers, 2019; Gonzales and Fuchs, 2017; Hsu et al., 2014). These tissues rely on resident tissue stem cells, which proliferate to self-renew and replenish the differentiated progeny of the epithelium. Within these tissues, self-renewal and differentiation must be tightly balanced and coordinated to prevent excessive growth or poor regeneration and healing. To tailor their activity, the stem cells reside in niches, where they communicate with their local microenvironment to receive the signals that regulate their maintenance and differentiation (Gehart and Clevers, 2019; Gonzales and Fuchs, 2017; Hsu et al., 2014).

The multifariousness of stem cell niches of barrier epithelia is exemplified by the intestine. The small intestine is organized into repeating crypt-villus units comprising a vast epithelial surface area and is the body’s primary site of absorption, whereas the large intestine is organized into repeating crypt units heavily colonized with beneficial bacteria and is a key site of vitamin and metabolite synthesis. As the body’s primary absorptive tissue, both, the small and large intestine, are subject to dynamic and systemic regulation, indicating a requirement for intestinal stem cells (ISCs) to integrate cues from their niches and coordinate stem cell behavior on a tissue-wide scale (Clevers, 2013). Since the initial description of ISCs within the crypt, however, it has become clear that these niches are diverse in their constituents and signaling pathways (Aoki et al., 2016; Baghdadi et al., 2021; Biton et al., 2018; Degirmenci et al., 2018; Greicius et al., 2018; Lindemans et al., 2015; McCarthy et al., 2020; Roulis and Flavell, 2016; Sato et al., 2011; Shoshkes-Carmel et al., 2018; Stzepourginski et al., 2017). While it has been surmised that key signals involved in ISC maintenance, such as WNTs and inhibitors of BMPs, may exist as gradients along the intestinal crypt-villus axis to guide intestinal differentiation (Beumer and Clevers, 2021; Gregorieff et al., 2005; He et al., 2004), the nature and sources of many of these signals still remain incompletely understood. Also awaiting investigation is how these signals might differ between the small intestine and less well studied large intestine, and how signals are coordinated across crypts to instruct stem cell behavior in the tissue.

Improvements in imaging techniques and spatial transcriptomics provide new opportunities to map the niche and identify dynamically responsive 3-dimensional (3D) features, such as the vasculature, with higher resolution than ever before. Such deep 3-dimensional imaging has provided insights into how tissue stem cells respond to their microenvironment, including through interactions with blood vessels and lymphatic capillaries (Comazzetto et al., 2021; Gur-Cohen et al., 2019). In the small intestine, lacteal capillary extensions of the submucosal lymphatic vasculature reach into the absorptive intestinal villi and participate in nutrient absorption and immune surveillance (Bernier-Latmani and Petrova, 2017). Although the large intestine lacks lacteals and the lymphatic structure in this tissue is less well characterized, submucosal lymphatic alterations in both the small and large intestine are an important pathologic feature in intestinal tissue inflammation in inflammatory bowel disease, a multifactorial disease characterized by inflammation and epithelial perturbations (Van Kruiningen and Colombel, 2008; Steven Alexander et al., 2010). Whether lymphatics regulate the ISCs of either the small or large intestinal epithelium remains unexplored and difficult to address given the spatial complexity of the lymphatic vasculature throughout these tissues. Here, we uncover lymphatic capillaries as hitherto unappreciated components of both small and large intestinal crypts, and we show that they act as key regulators of intestinal epithelial differentiation. Employing whole tissue clearing and 3D imaging, we reveal an intimate association between the lymphatic capillary network and the base of intestinal crypts, where ISCs reside. By devising a lymphatic endothelial cell (LEC):organoid coculture system, we recapitulate this association *in vitro* and show that lymphangiocrine, i.e. LEC-secreted, factors alter intestinal organoid morphogenesis and maturation.

To delineate the nature of the putative LEC:ISC crosstalk, we perform single cell RNA-sequencing (scRNA-seq) and spatial transcriptomics on both, the small intestine (replete with lacteals) and large intestine (lacking lacteals) of adult mice. Devising a new computational approach, we deconvolute these low-resolution spatial transcriptomic profiles to reconstruct the spatial relationships of the diverse intestinal cell types along the stereotypical crypt-villus and crypt axes. The ability to assign cell type-specific transcriptomes according to their cellular location along this axis has enabled us to cartograph the cellular and transcriptional landscape of these tissues at high resolution. In doing so, we identify a unique lymphatic regulatory program at the crypt base that is distinct from lymphatic functions within the villi. By combining these newfound computational methods with functional interrogation, we unearth spatial ISC receptor-lymphatic ligand interactions and their functional importance in regulating ISC behavior and governing tissue dynamics.

## Results

### Lymphatic capillaries neighbor stem cells at the intestinal crypt base

The cellular niche of ISCs residing within the pocket-like crypts of the small intestinal epithelium includes neighboring specialized epithelial cells (Paneth cells), sub-epithelial mesenchymal cells (pericryptal fibroblasts, myofibroblasts, stromal telocytes, trophocytes) and immune cells (Zhu et al., 2021). Intestinal stem cell identity is lost as committed short-lived proliferative progeny continue their upward journey along the crypt-villus axis and then differentiate into one of the different secretory and absorptive lineage options according to their microenvironment, nutrient availability and inflammatory cues (Beumer and Clevers, 2021). In normal homeostasis when environmental cues are largely stable, stem cells within individual crypts regenerate the epithelium at largely steady rates.

Recently, we discovered that lymphatic capillaries are tightly associated with hair follicle stem cells (HFSCs), where they synchronize hair regenerative activity across the skin and govern quiescence of individual hair follicle niches (Gur-Cohen et al., 2019). Compared to HFSCs, which undergo long bouts of quiescence, however, ISCs proliferate continuously, leading us to wonder whether in the intestine, another epithelial barrier tissue, lymphatic capillaries would still possess a stem cell regulatory function, and if so, how.

As vascular networks like lymphatics are challenging to visualize by conventional imaging analyses, we set out to characterize the cellular diversity surrounding the intestinal stem cell niche, applying tissue clearing and whole-mount imaging, which allows for visualization of the 3D landscape of the ISC niche, including the complex architecture of the vasculature (Richardson and Lichtman, 2015).

Our 3D-imaging of the small intestine revealed the well-known capillary plexus composed of a blood-vascular network that surrounds a central lymphatic capillary (lacteal) along the villus axis (Figure S1A and S1B and Video S1) (Bernier-Latmani and Petrova, 2017). As prominent as lacteals, however, and often overlooked, was a rich vascular network of lymphatic capillaries residing just beneath the crypt. Co-immunofluorescence of the lymphatic marker LYVE1 and ISC marker LGR5^+^ exposed an intriguing association of these lymphatic capillaries and ISCs, spanning from one crypt base to another (Figures 1A, S1A and S1B and Video S2).

**Figure 1.**
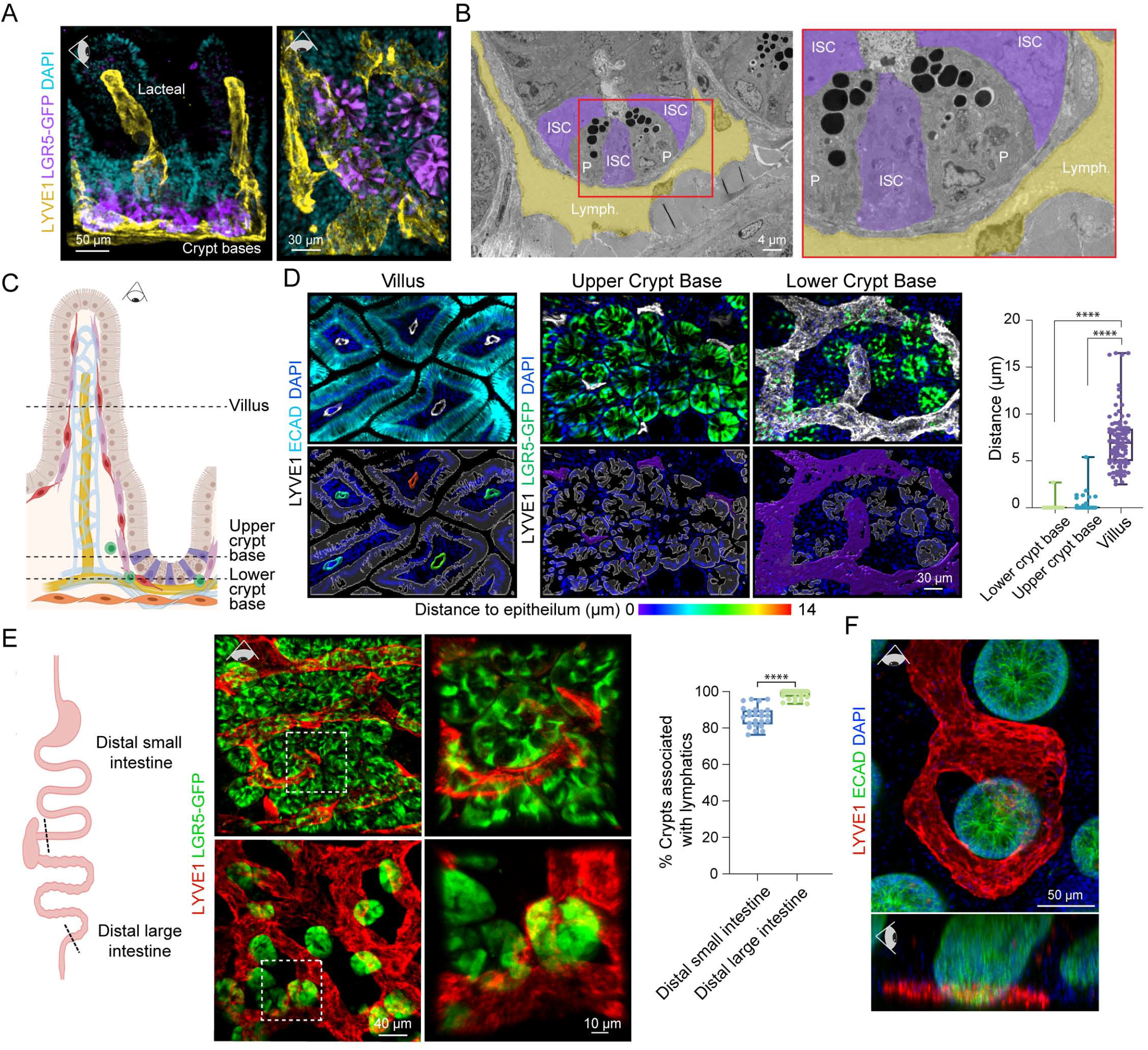
Lymphatic capillaries nest crypt-based intestinal stem cells. **(A)** 3D immunofluorescence of small intestinal crypts and villi structures, revealing LYVE1^+^ lymphatic capillaries (yellow) nesting LGR5^+^ crypt-based stem cells (purple). Representative of *n >* 5 mice. Eye icon indicates the plane of view of visualization; horizontal eye, side view; downward facing eye, top view with tissue lying flat. **(B)** Pseudocolored Transmission Electron Microscopy images of the small intestinal crypt base with lymphatic capillaries (yellow, Lymph.), Paneth cells (P), and intestinal stem cells (purple, ISCs). Red box inset is magnified in the right side panel. **(C)** Schematic of the small intestinal crypt niche, showing ISCs (purple) intermingled with Paneth cells (legend in Figure S1A). **(D)** 3D immunofluorescence images of 5 µm z-stacks of the mouse small intestinal lymphatic vasculature and epithelial cells (villus) or ISCs (upper and lower crypt base) as shown in the schematic (dotted lines) in C. Quantification of the distance between vascular surfaces and the epithelium was done in each of the three regions (*n =* 9 mice with ≥ 3 images/region/mouse). **** indicates a p-value of <0.0001 (One-way analysis of variance (ANOVA), Tukey’s multiple comparisons) **(E)** 3D immunofluorescence images of cleared small (top) and large (bottom) intestine, demonstrating the percentage of LGR5^+^ crypts (green) that are associated with lymphatic capillaries (red) with quantification (*n =* 9 mice with ≥ 4 images/region/mouse). Boxed images are magnified at the right. *** indicates a p-value of 0.0001 to 0.001 (Unpaired two-tailed Student’s *t*-test) **(F)** Representative human intestinal biopsy after tissue clearing and 3D imaging, demonstrating conserved association between lymphatics (LYVE1^+^, red) and the intestinal crypt base (ECAD^+^, green) in the distal colon (representative of *n =* 4 individual patients). All scale bars appear on each image; antibodies are color-coded.

Ultrastructural analysis revealed the intimate physical proximity between lymphatics and crypt base-resident ISCs, often separated only by a thin layer of extracellular matrix (ECM) (Figure 1B). Markers against previously identified mesenchymal and immune cells in the crypt revealed that the lymphatic network was interwoven with the components of the ISC niche (Figures 1C and S1C-S1F). In some cases, lymphatic capillary protrusions were found in direct contact with the crypt base (Figure S1G).

Lymphatic lacteals are well-established pathways for nutrient absorption and immune cell trafficking in the intestine (Cifarelli and Eichmann, 2019). In the villus, lacteals were separated from the epithelium by a network of blood vessels, immune cells, and abundant stromal cells (Figures S1A-S1H). By contrast, lymphatic capillaries at the crypt base were closely associated with resident epithelial cells, including Paneth and LGR5^+^ stem and progenitor cells (Figures 1A and 1D).

Given the critical function of small intestinal lymphatics in nutrient absorption, we considered that lymphatic endothelial cell proximity to the crypt base might be merely a necessary component of their path from the lacteals to the collecting vessels which flow into mesenteric lymph nodes and ultimately deliver nutrients into the systemic circulation through the thoracic duct (Bernier-Latmani and Petrova, 2017). If lymphatic:SC proximity in the small intestine is simply a consequence of lacteal collection then we would not expect to observe this in the large intestine, as the lymphatic vascular system of the large intestine is devoid of lacteals and plays a nominal role in nutrient absorption (Bernier-Latmani and Petrova, 2017).

Colonic lymphatics, i.e. lymphatics of the large intestine, are known to have important roles in intestinal immune surveillance and regulation (Esterházy et al., 2019; Houston et al., 2015), yet the structure of the capillary network has not been fully characterized. Intriguingly, crypt-based LGR5^+^ SCs were associated with lymphatic capillaries across the intestine (Figure 1E). While the colonic ISC niche lacks Paneth cells, the majority of LGR5^+^ crypts in the large intestine were positioned at least partially atop lymphatic capillaries (Figures 1E, S2A-S2E and Video S3), reminiscent of what we observed in the small intestine.

Ultrastructural analysis revealed a similar close proximity between ISCs and lymphatic capillaries in the large intestine (Figure S2C). Notably, the lymphatic-ISC association was also conserved in humans, where we identified lymphatic capillaries in association with crypt-based epithelial cells in large intestinal biopsy specimens (Figure 1F and Video S4). When taken together with our prior findings on the lymphatic capillary network of the hair follicle, these data establish that lymphatic capillaries can form intricate associations with epithelial stem cell niches irrespective of whether the stem cells are quiescent (hair follicle) or rapidly cycling (large and small intestines). These new revelations begged the question as to whether there might be a common purpose of this close encounter and a role for lymphatics in tissue maintenance. Thus, we sought to define and characterize the phenotypic impact of lymphatic capillaries on ISC behavior.

### Lymphatic endothelial cells instruct intestinal organoid development and maturation

To test whether lymphatic endothelial cells signal directly to the intestinal epithelium, we took advantage of the ability to culture small intestinal epithelial stem cells to generate 3D “mini-gut” organoids (Sato et al., 2009) and established a system to coculture organoids with primary murine LECs (Figure 2A). This system recapitulated the connection between lymphatic endothelial cells and small intestinal stem cells observed *in vivo* and enabled us to distinguish between effects due to direct cell-to-cell communication and those resulting from lymphatic drainage properties that are absent in this coculture system (Figures 2A, S3A-S3B and Videos S5-S6). Under coculture conditions (50% conventional organoid media:50% LEC media), LECs maintained expression of key lymphatic transcription factors and protein markers (PROX1, CD31 and LYVE1) as well as their ability to form endothelial tubes when cultured on Matrigel in this medium (Figure S3C). Similarly, organoids grown alone in coculture medium also maintained key features, including the ability to form crypt domains (Figure S3D).

**Figure 2.**
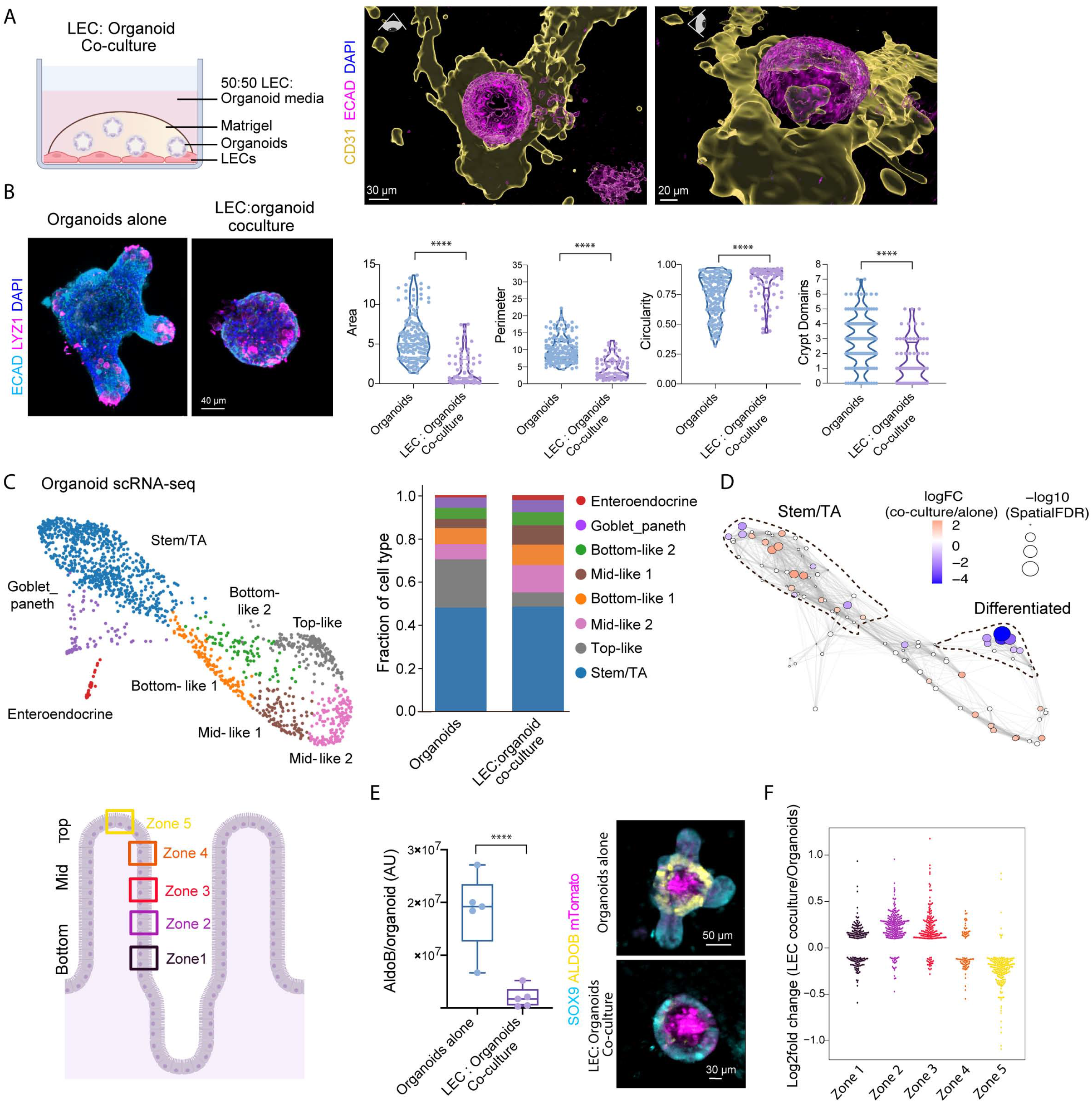
Lymphatic endothelial cells instruct intestinal organoid development and maturation. **(A)** Illustration of the LEC:organoid coculture system (left panel). Organoids were grown either alone or on top of LECs in 3D Matrigel and 50% LEC:50% organoid (ENR) media. Right panel: Representative 3D image reconstruction of an organoid (ECAD^+^, pink) associated with LECs (CD31^+^, yellow) in culture. **(B)** Phenotypic characterization of organoids cocultured with LECs. Representative image of an organoid (ECAD^+^, light blue) and Paneth cells (LYZ1^+^, magenta), and the respective quantification of organoid size and morphology (area, perimeter, circularity and crypt domains). **** indicates a p-value of <0.0001 (Unpaired two-tailed Student’s *t*-tests). **(C)** Force-directed layout generated from scRNA-seq data of organoids cultured in the presence or absence of LECs. Each cell is colored according to cell type and then contrasted according to representation within each culture condition. Relative distribution of cell types per condition is indicated in the bar plot in the middle. Intestinal villus zones used to generate the comprehensive spatial clustering of enterocytes are shown in the schematic on the right (Moor et al., 2018). **(D)** Nearest-neighbor differential abundance analysis of cell states in the scRNA-seq data was based on variation of the Milo algorithm (Dann et al., 2021). In this analysis, nodes are neighborhoods, colored by their log_2_ fold change of cell abundance in LEC-cocultured organoids vs. organoids alone, and sizes correspond to the minus log_10_ false discovery rate (FDR). The position of nodes is determined by the position of the neighborhood index cell from the force-directed layout (shown in C). Stem/progenitors and differentiated enterocytes are outlined as shown. **(E)** Organoids derived from an mTmG mouse (mTomato, magenta) were grown alone or in coculture with LECs, and imaged (maximum intensity projections) by immunolabeling for SOX9^+^ progenitors (light blue) and Aldolase B differentiated enterocytes (ALDOB^+^; yellow). Quantification of ALDOB signal intensity is at left. *n =* 3 independent experiments. ** indicates a p-value of 0.001 to 0.01 (Unpaired two-tailed Student’s *t*-test) **(F)** Log_2_ fold change of the transcription levels of villus enterocyte zone signature genes (Moor et al., 2018) in organoids cocultured with LECs vs. organoids grown alone.

ISCs or isolated crypts cultured together with lymphatic endothelial tubes generated organoids with significant morphologic changes compared to crypts cultured on their own. Organoids formed in the presence of LECs were smaller in size, more circular in shape and contained fewer crypt domains, suggestive of an impaired maturation (Figures 2B, S3B and S3E). Importantly, however, these organoids displayed lysozyme-producing Paneth cells, a critical component of the small intestinal crypt and essential for ISC maintenance (Figure 2B). This, and the maintained ability of organoids to form in the presence of LECs, suggested that lymphatic capillaries did not impair organoid formation *per se*. Rather, the presence of LECs appeared to prevent their maturation. This effect was not observed when ISCs were cocultured with blood vascular endothelial cells (BECs), underscoring the specificity of the lymphatic endothelium in repressing crypt differentiation (Figure S3F).

To gain further insights into the nature of these effects, we performed single cell RNA-sequencing (scRNA-seq) on organoids derived from LEC-coculture and compared them to scRNA-seq profiles that we derived from standard organoid cultures grown in the same media. We pooled the cells from the two culture conditions and performed clustering (Levine et al., 2015). Cell types clustered independently of culture conditions (Figure S3G) and segregated into 11 clusters (Figure S3H). Two clusters expressing markers for Goblet/Paneth cells (e.g. *Lyz1*, *Muc2*) and enteroendocrine (e.g. *Chga*, *Chgb*) were assigned unambiguously (Haber et al., 2017). Additionally, four clusters expressing markers for both stem/progenitor and transit-amplifying (TA) cells (e.g. *Olfm4*, *Ascl2*, *Tubb5*, *Top2a*) were labeled as stem/TA (Haber et al., 2017; Nikolaev et al., 2020). Lastly, five clusters expressed enterocyte markers, such as *Alpi* and *Fabp1*, and correlated with enterocyte differentiation programs along the axis of villus zone positions (Moor et al., 2018) (Figure 2C, bottom); we labeled them accordingly as bottom-like 1, bottom-like 2, mid-like 1, mid-like 2, top-like (Figures 2C, S3H and S3I).

Organoids grown in the presence of LECs displayed substantially fewer terminally differentiated enterocytes though similar proportions of stem and progenitor cells (Figure 2C). These findings were also concordant with a nearest-neighbor graph-based differential abundance analysis (Milo) (Dann et al., 2021), which provided us with a finer resolution (Figure 2D). This analysis yielded further insights, revealing that organoids grown in the presence of LECs displayed altered distributions of cells within the stem/progenitor cluster and a skewing of enterocyte progenitors at the expense of fully (terminally) differentiated enterocytes (Figure 2D).

LEC-cocultured organoids were also underrepresented in the cell fraction corresponding to actively cycling (*Mki67^+^)* cells (Figures 2D and S3J). Consistent with the reduced population of actively cycling intestinal stem/progenitor cells, incorporation of 5-ethynyl-20-deoxyuridine (EdU) during a 10 minute-pulse was reduced in organoids grown in the presence of LECs (Figure S3K). Consistent with a block in enterocyte terminal differentiation was the diminished immunofluorescence for the mature enterocyte marker, Aldolase B (ALDOB) (Figure 2E). Additionally, at the transcriptomic level, enterocytes from the co-cultured condition up-regulated genes defining bottom zones compared to those defining top zones (Figure 2F).

To better understand the difference in maturation between the two conditions, we computationally modeled the differentiation trajectory and quantified likely paths of differentiation (STAR methods). Enterocytes obtained from control organoids were more frequent within the terminally differentiated top zone, whereas those from LEC-cocultured organoids were more evenly distributed across intermediate states (Figure S3L). Taken together, these data support a model whereby lymphatic endothelial cells at the crypt base maintain stem and progenitor cells but restrict their differentiation into mature enterocytes.

### The lymphatic secretome maintains ISCs but restricts their terminal differentiation

If as our studies thus far predict, crypt lymphatics function by supporting the maintenance of intestinal stem and progenitor cells over rapidly cycling and terminally differentiated cells, then lymphatic endothelial cells should enhance the efficiency of forming secondary organoids from single cells dissociated from a primary (naïve) organoid. Further, if crypt lymphatics also function to prevent the terminal differentiation of ISCs into mature enterocytes, then the size of organoids emerging in this assay should be reduced compared to secondary organoids formed in the same culture media but in the absence of LECs.

To test these hypotheses, we first assessed the capacity of LECs to support organoid growth. Indeed, the presence of LECs more than doubled the organoid forming efficiency of single ISCs (Figure 3A). Similar to crypt-derived organoids, single cell-derived organoids grown in the presence of LECs were smaller with fewer protrusions. As predicted, LEC-cocultured organoids also exhibited decreased ALDOB immunofluorescence, consistent with a block in the generation of enterocytes (Figure 3B).

**Figure 3:**
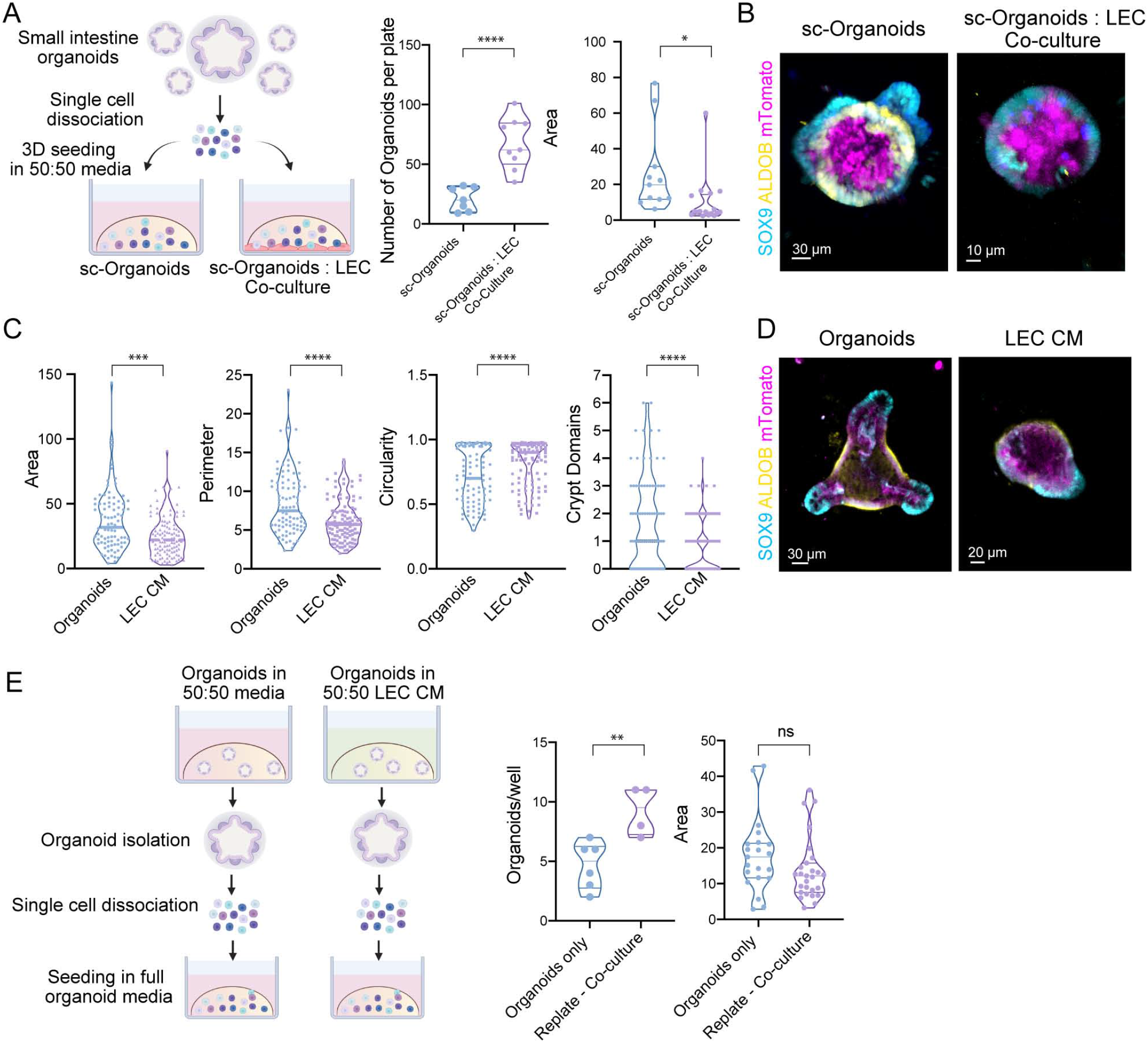
Lymphatic signals maintain organoid-generating capacity and reduce terminal differentiation of enterocytes. **(A)** Schematic of the single-cell assay. Single cells from primary organoids were seeded in 50% organoid:50% LEC media in the presence or absence of LECs. Quantifications of organoid number and area are on the right. Data are from *n =* 3 independent experiments with ≥ 2 wells/condition. **** indicates a p-value of <0.0001, * indicates a p-value of 0.01 to 0.05 (Unpaired two-tailed Student’s *t*-test). **(B)** Aldolase B (ALDOB) expression in single-cell-derived organoids grown as shown in A. Images are maximum intensity projections showing organoids derived from an mTmG mouse (mTomato, magenta) and stained for SOX9 (light blue) and ALDOB (yellow). **(C)** Crypt-derived organoids were cultured alone in 50:50 control and LEC-conditioned media. Violin plots of the area, perimeter, circularity and number of crypt domains are shown following culture for 4 days. **** indicates a p-value <0.0001, *** indicates a p-value of 0.0001 to 0.001 (Unpaired two-tailed Student’s *t*-test) **(D)** Immunofluorescence staining as in B of crypt-derived organoids grown in 50:50 control or LEC-conditioned media. **(E)** Schematic of the replate assay. Single cells derived from organoids grown in 50:50 control or LEC-conditioned media were re-seeded in 100% organoid (ENR) media. Quantification of organoid number and area, measured in *n =* 3 independent experiments with ≥ 2 wells per condition, is shown on the right. ** indicates a p-value of 0.001 to 0.01, ns means non-significant (Unpaired two-tailed Student’s *t*-test).

Not all organoids made contact with LECs, yet still most displayed reduced size and differentiation (Figure S3B). Similarly, many but not all crypts showed close association with lymphatic capillaries *in vivo* and even when they did so, direct cell-to-cell contact between LGR5^+^ stem cells and LECs appeared to be quite rare (Figure 1E). We therefore reasoned that LEC-secreted (lymphangiocrine) factors might be responsible for the phenotype observed in cocultured organoids. To test this hypothesis, we collected LEC-conditioned medium and examined its potential to affect organoid initiation and maturation in the absence of LECs themselves. LEC-conditioned medium recapitulated the phenotype of LEC-cocultured organoids, displaying smaller organoids with less differentiated progeny (Figures 3C and 3D).

We next isolated single cells from organoids grown in LEC-conditioned medium and then plated them in normal organoid culture conditions without LECs (Figure 3E). Two important findings emerged. First, LEC-conditioned media enhanced the organoid colony-forming efficiency of LEC-conditioned ISC progenitors in these secondary assays (Figure 3E), thereby underscoring the potent and direct effect of lymphangiocrine factors on ISC maintenance. Second, the organoids that formed from these pre-conditioned ISCs differentiated normally once the lymphangiocrine factors were no longer present (Figure 3E), indicating that the effects of LECs in impairing ISC differentiation are reversible and dependent upon sustained exposure to these lymphangiocrine factors. In the context of the tissue, such effects should be manifested predominantly within the crypt niche, where the stem cells and lymphatics reside and where their differentiation is suppressed.

### The *in vivo* spatial map of cell types in the small and large intestines

We next turned to addressing the nature of the crypt lymphangiocrine factors and unraveling how they impact intestinal stem cells *in vivo.* Since lacteal lymphatics might produce an entirely different set of factors than lymphatics located at the base of the crypt, meaningful results necessitated a spatial transcriptomic approach. Therefore, we sought to cartograph intestinal cell types and their gene expression profiles along the spatial dimensions of the crypt-villus axis (for small intestine) and crypt axis (for large intestine) by computationally integrating scRNA-seq and spatial transcriptomic data (Figure 4A). This approach allowed us to (1) identify genes highly and/or uniquely expressed by LECs and (2) correlate LEC gene expression patterns with the spatial proximity between lymphatics and ISCs (or other potential receiver cells) within the intestinal epithelium.

**Figure 4.**
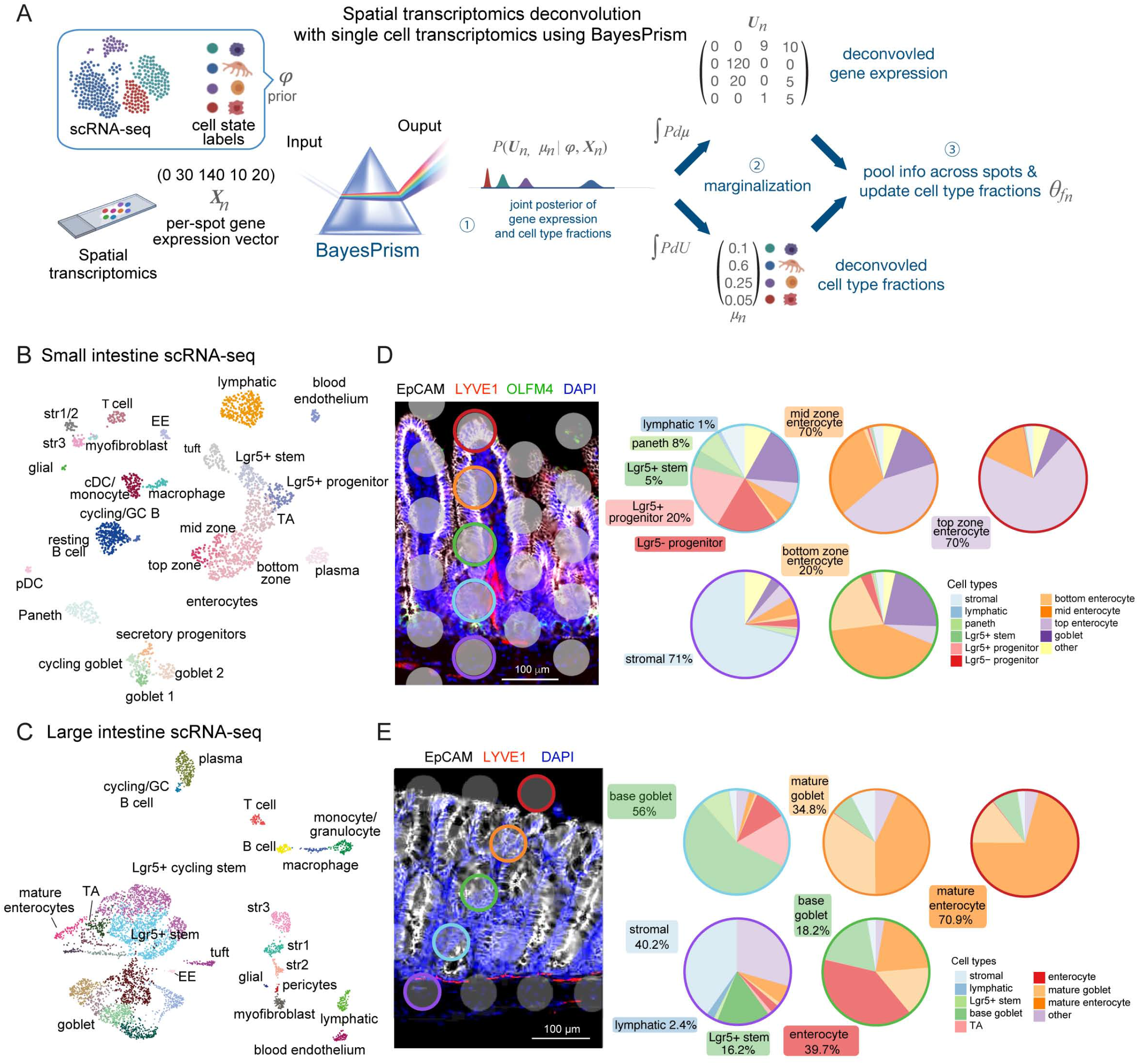
Integrated transcriptomics reconstructs the cellular and molecular landscape of the crypt-villus axis. **(A)** Workflow of spatial transcriptomic data analysis. The left panel shows the algorithmic workflow of single-cell:spatial transcriptomic data integration and deconvolution based on BayesPrism to infer joint gene expression and cell type fraction per spot of the 10X Visium gene expression slide layout. (B and C) Uniform Manifold Approximation and Projection (UMAP) plots of murine scRNA-seq data of small and large intestinal tissue, respectively. Each cell is colored according to cell type annotation. EE: enteroendocrine cell, pDC: plasmacytoid dendritic cell, cDC: classical dendritic cell, str: stromal cell and TA: transit-amplifying cell. Data are enriched for lymphatic endothelial cells and LGR5^+^ stem cells. (**D** and **E**) Left: Representative immunofluorescence images from the small and large intestinal tissue sections, respectively, used for spatial transcriptomic (10X Visium) profiling. The gray dots each correspond to the 50 µm diameter areas captured for RNA-seq. The overlaid immunofluorescence marks EpCAM^+^ epithelial cells (white), LYVE1^+^ lymphatic vasculature (red), OLFM4^+^ intestinal stem cells (green) and DAPI^+^ nuclei (blue). Right: Pie charts represent the fractions of selected cell types of interest in each correspondingly colored spot along the crypt-villus axis deconvolved by BayesPrism. For visualization, multiple crypt-based goblet subtypes were grouped and labeled as base goblet.

We collected comprehensive scRNA-seq profiles of intestinal cells from both small and large intestine, covering the major cell types from immune, stromal and epithelial lineages within the intestine, while enriching for rare populations of LECs and LGR5^+^ ISCs (Figures 4B, 4C, and S4A-S4E). Immune cells of the small and large intestine each encompassed the major lymphoid (T cells, B cells, plasma cells and germinal center B cells) and myeloid (macrophages, dendritic cells, monocytes and granulocytes) cell populations. Analogously, stromal cells clustered according to glial (e.g. *S100b*, *Gfap*), blood endothelial (e.g. *Cd31*, *Cdh5*) and lymphatic endothelial (e.g. *Lyve1*, *Prox1*), myofibroblast (e.g. *Acta2*, *Myh11*) and fibroblast (e.g*. Col6a2, Dpt*) marker genes (Kinchen et al., 2018) (Figure 4B, 4C, S4A and S4B). Our clustering further revealed heterogeneity in mesenchymal cell populations as defined by markers for trophocytes (e.g. *Cd81*), telocytes (e.g. *Foxl1*) and pericryptal stromal cells (e.g. *Cd34, Pdpn*) (Figures 4B, 4C, S4C and S4D). Finally, epithelial cells of the small and large intestine reflected all of the main differentiated lineages as highlighted by markers for enterocytes (e.g. *Alpi*, *Fabp1*), goblet cells (e.g. *Muc2*), enteroendocrine cells (e.g. *Chga, Chgb*), tuft cells (e.g*. Dclk1*) and Paneth cells (e.g*. Lyz1, Defa17*), with the latter being absent in the large intestine (Figures 4B, 4C, S4A and S4B). Goblet cells exhibited a high degree of heterogeneity and accounted for a large proportion of epithelial cells in the large intestine. Immature epithelial cells segregated into secretory precursors (e.g*. Atoh1, Dll1*), transit-amplifying cells (e.g. *Stmn1, Tubb5*) and Lgr5^+^ progenitors. *Bona fide* ISCs marked by *Lgr5* and *Olfm4* were found in both scRNA-seq datasets.

While these early populations only comprise 4-6% of the total crypt epithelial cells in the small intestine, our enrichment strategy allowed us to increase the amount of LGR5^+^ stem/progenitor cells to ∼30% of the total population, providing a deep molecular signature of these rare populations (Barker et al., 2012). Taken together, these data constitute an unprecedented comprehensive atlas of mouse colonic epithelium, and a valuable resource of transcriptomes for mouse small and large intestinal cell populations in general and ISCs in particular.

Digging deeper, we next performed spatial transcriptomics (10X Visium platform) on matched tissue samples that were immunolabeled for epithelial cells (EpCAM), ISCs (OLFM4) and LECs (LYVE1) (Figures 4D, 4E, S5A and S5C). Commercially available platforms for spatial transcriptomics (10X Visium) are a valuable tool, yet limited in resolution, with each barcoded spot (∼ 50 µm diameter spots with a 100 µm center to center distance) typically containing multiple cells and cell types. Such limitations precluded the characterization of interactions between adjacent cells. This was particularly problematic for our interest, namely to dissect the interactions between ISCs and their niche components, as these cell types were often combined within a single spot. Understanding the nature of the interactions between cell types in the niche necessitated methods to deconvolute (1) the cell types within each spot and (2) the genes that each cell expresses at a given spatial position.

For this purpose, we used BayesPrism, a Bayesian statistical model that jointly deconvolves cell type composition and cell type-specific gene expression profiles within each spatial spot by using a scRNA-seq reference from matched tissue as prior information (Figure 4A) (Chu et al., 2021; McKellar et al., 2020). This method significantly outperforms other regression-based tools in bulk RNA-seq deconvolution (Chu et al., 2021; McKellar et al., 2021) (Figure S5A), and its robustness to platform batch effects, technical artefacts and noise made it particularly well suited for spatial deconvolution, treating each Visium spot as a bulk RNA sample. Moreover, our enrichment for LECs and LGR5^+^ ISCs in our scRNA-seq strategy allowed for an accurate inference of these rarer cell types across the intestine (Figures S5B and S5C).

We validated our deconvolution assignments by comparing them to the expected cell type distribution along the crypt-villus or crypt axes and to immunofluorescence profiles of the sequenced tissue (Figures 4D-4E). Benchmarking BayesPrism against other deconvolution tools developed for spatial transcriptomic data, we found that BayesPrism showed the highest concordance with expected marker gene expression patterns (Figures S5A-S5C and STAR methods) (Cable et al., 2021; Kleshchevnikov et al., 2020; Lopez et al., 2021; McKellar et al., 2021).

### Cell type and expression cartography reveals *in vivo* lymphatic:ISC interactome

While Visium allows charting the spatial patterns of gene expression, its limited resolution and sparseness in gene measurements precluded the high-resolution spatial gene expression maps we needed to interrogate niche: ISC interactions (Figures 4D, 4E, S5A and S5C). To build a high-resolution cartograph of gene expression along the crypt-villus axis capable of revealing cell-cell interactions, we developed a SpaceFold axis projection (Figure 5A). For this purpose, we used non-linear dimensionality reduction of cell-type fractions per spot to project each Visium spot along a 1-dimensional (1D) pseudo-space axis.

**Figure 5.**
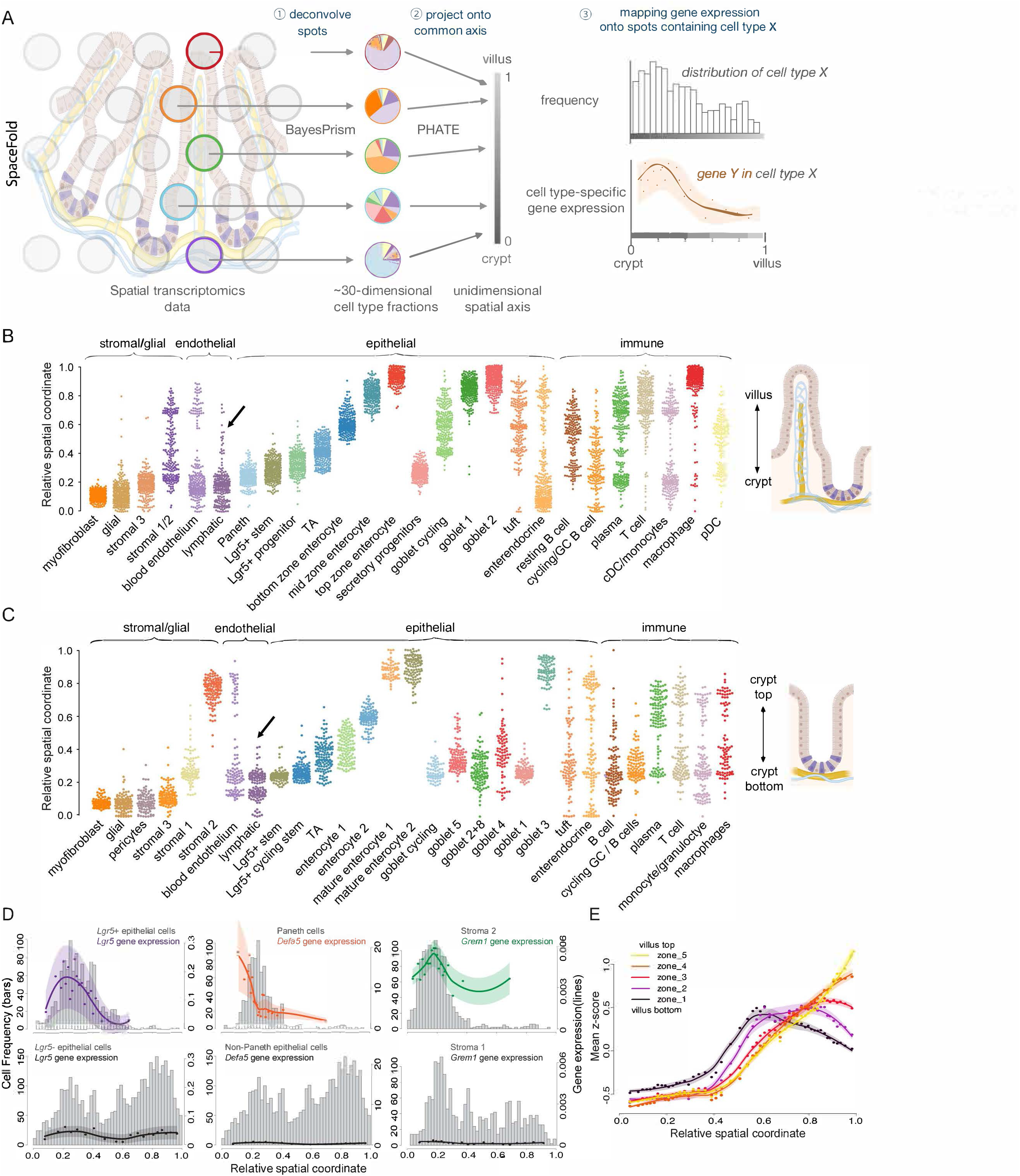
SpaceFold cartography reveals the lymphatic:ISC interactome in spatial resolution. **(A)** Workflow of SpaceFold, illustrated using crypt-villus units. Each spatial spot was deconvolved using BayesPrism (step 1) to generate information about cell type fractions. Vectors of cell type fractions were reduced to a 1-dimensional projection that approximates its physical position along the crypt or crypt-villus axis using PHATE (Moon et al., 2019) (step 2). BayesPrism’s cell type-specific gene expression was used to generate trends along this axis for spots containing the cell type(s) of interest (step 3). **(B-C)** SpaceFold reveals the relative spatial coordinates of cell types, as identified from individual spots, along the small intestinal crypt-villus axis (B) and large intestinal crypt axis (C). Cell type annotation follows the cluster nomenclature in figures 4B-4C. Each dot represents a Visium spot which contains the indicated cell-type, plotted along its SpaceFold projection. The black arrow in B denotes computationally reconstructed lymphatic lacteals in the small intestine, which are missing in the large intestine in C. **(D)** SpaceFold maps cell type-specific expression of known cell type markers onto the crypt-villus axis in the small intestine. X-axes mark the relative spatial coordinate imputed by SpaceFold. Histograms show the distribution of spots containing the selected cell types along the crypt-villus axis. Y-axes on the left indicate the frequency of spots containing the cell type (related to the histogram). Y-axes on the right indicate the expression level of the indicated gene in the indicated cell-type (related to the dots, lines and shaded areas). For comparison between a group of cell types (left and center panels), y-axes on the right indicate raw expression values, *U̅*, inferred by BayesPrism summed over cell types within the selected group. For comparison between individual cell types (right panels), y-axes on the right indicate the expression value *U* normalized by the total amount of transcription in the cell type of interest (see Methods for details). Each dot represents the mean expression over an interval of selected spots binned along the SpaceFold axis (x-axis). The x-coordinate of each spot represents the mean of the SpaceFold coordinates in each bin. Lines mark the mean values fitted using local polynomial regression. Shaded areas represent the mean ± 2 standard error. Top panels show the predicted expression in cell types expected to express the indicated marker genes, while bottom panels show the predicted expression in cell types not expected to express those genes. (**E**) Intestinal villus epithelial cell zone gene expression distributed spatially along the projected crypt-villus axis. Similar to **D**, mean z-scores of the library size-normalized expression of each group of zone markers in enterocytes were plotted on the SpaceFold crypt-villus axis. Each dot represents the mean z-scores averaged over an interval of all spatial spots binned by the SpaceFold coordinate. The x-coordinate of each spot represents the mean of the SpaceFold spatial coordinates in each bin. Lines mark the mean values fitted using local polynomial regression. Shaded areas represent the mean ± 2 standard error.

SpaceFold took advantage of the hundreds of highly stereotyped structures of the intestinal epithelium, each comprised of repeating crypt-villus units in the small intestine, or crypt units in the large intestine. Despite inherent differences between the small and large intestine, each crypt within the respective tissues is similar in size and ordering of cell types along the crypt-villus or crypt axis (Figure 5A). We hypothesized that, as spatial spots were randomly sampled along this stereotypical axis, the cell type composition of each spatial spot, as inferred by BayesPrism, would inform the relative physical coordinates of cells along a 1D artificial crypt-villus unit (STAR methods).

Indeed, one-dimensional pseudospace inferred by SpaceFold beautifully recapitulated the expected cell type distributions along the small intestinal crypt-villus and large intestinal crypt axes, respectively (Figures 5B and 5C). Absorptive enterocyte clusters mapped along the villus axis as predicted by villus zone marker genes, with bottom-and top-zone enterocytes falling into their respective positions. Interestingly, the projected axis distinguished the spots containing the crypt-based lymphatic capillaries from the lacteal lymphatics, which spanned from the base lymphatic capillary bed to the tips of the intestinal villi in the small intestine (Figure 5B, arrow). Notably, such protrusions were not present in the large intestine data set, consistent with the absence of lacteals in this tissue (Figure 5C, arrow). Importantly, this cartography also recapitulated the known distribution of ISC niche cells along this axis, with the stromal 3-like population containing trophocytes falling in its predicted position below the crypt base in small and large intestine (Figures 5B and 5C) (Kinchen et al., 2018; McCarthy et al., 2020).

We next cartographed gene expression profiles for each cell type. Based on deconvolved cell type-specific gene expression profiles in each spatial spot, we mapped transcript levels of established epithelial (e.g. *Lgr5* as a marker of ISCs*, Defa5* as a marker of Paneth cells) and stromal (e.g. *Grem1* as a marker of crypt-based stromal cells/trophocytes) marker genes of each spot along our projected crypt-villus axis. A high concordance was found between marker gene expression and the expected spatial distribution of their associated cell types (Figure 5D). Intestinal villus epithelial cell zone markers also distributed predictably along the projected villus axis (Moor et al., 2018) (Figure 5E). Taken together, by accurately deconvolving the cell type composition, inferring cell type-specific gene expression profiles within each spatial spot, and generating a stereotypical axis projection via SpaceFold, we were able to derive a high-resolution cartograph of cell types and cell type-specific gene expression profiles in the small and large intestine, along the crypt-villus and crypt axes, respectively. Although the focus of our study was on LEC:ISC interactions in the crypt niche, our data and cartography provide a rich resource for charting interactions between other cell types in the intestine.

### Lymphatic capillaries are a local source of stem cell niche factors

With the deconvolved spatial transcriptome of the mouse intestine in hand and the knowledge that lymphatic-derived secreted factors modulate organoid behavior *in vitro* (Figures 3C-3E), we were poised to unravel the spatially relevant crypt lymphangiogenic ligands capable of signaling to ISCs *in vivo*. Genes expressed at significant levels in lymphatic endothelial cells (expressed in more than 20% of LECs), but not in related blood endothelial cells (expressed in less than 20% of BECs), were obtained from our single cell RNA-seq dataset. From these, we generated a list of 12 genes encoding proteins with a high probability of being secreted extracellularly based on the presence of sequences encoding N-terminal signal peptides that predict secretion and extracellular localization (Emanuelsson et al., 2000; Horton et al., 2007) (Figure S6A).

We next examined these 12 LEC-derived factors for their median expression among the different cell types of the crypt niche (Figure S6A). We excluded those with low probability of being directly relevant to ISC function (e.g. those encoding proteins with roles in antimicrobial peptides or proteins involved in clotting cascades or copper metabolism) to focus on a list of 5 candidate lymphangiocrine factors (Figure 6A). In parallel, we used our high-resolution, cell-type specific, pseudo-space transcriptomic cartograph to plot the spatial expression of each of these genes along the crypt-villus axis in a cell type-specific manner (Figure 6B). From these analyses, we learned that some of these genes, e.g. *Ntn1* and *Il33*, showed enrichment but not exclusive expression in lymphatics. By contrast, *Reln, Ccl21a, Rspo3* and *Wnt2,* were more highly expressed in LECs than other niche cells and additionally were enriched in the niche lymphatics at the crypt base. Of these four factors, expressed predominantly by crypt-based lymphatic endothelial cells, three had receptors that were expressed by ISCs in the small and large intestine, making them strong candidates for mediating LEC:ISC signaling within the crypt niche (Figure S6B).

**Figure 6.**
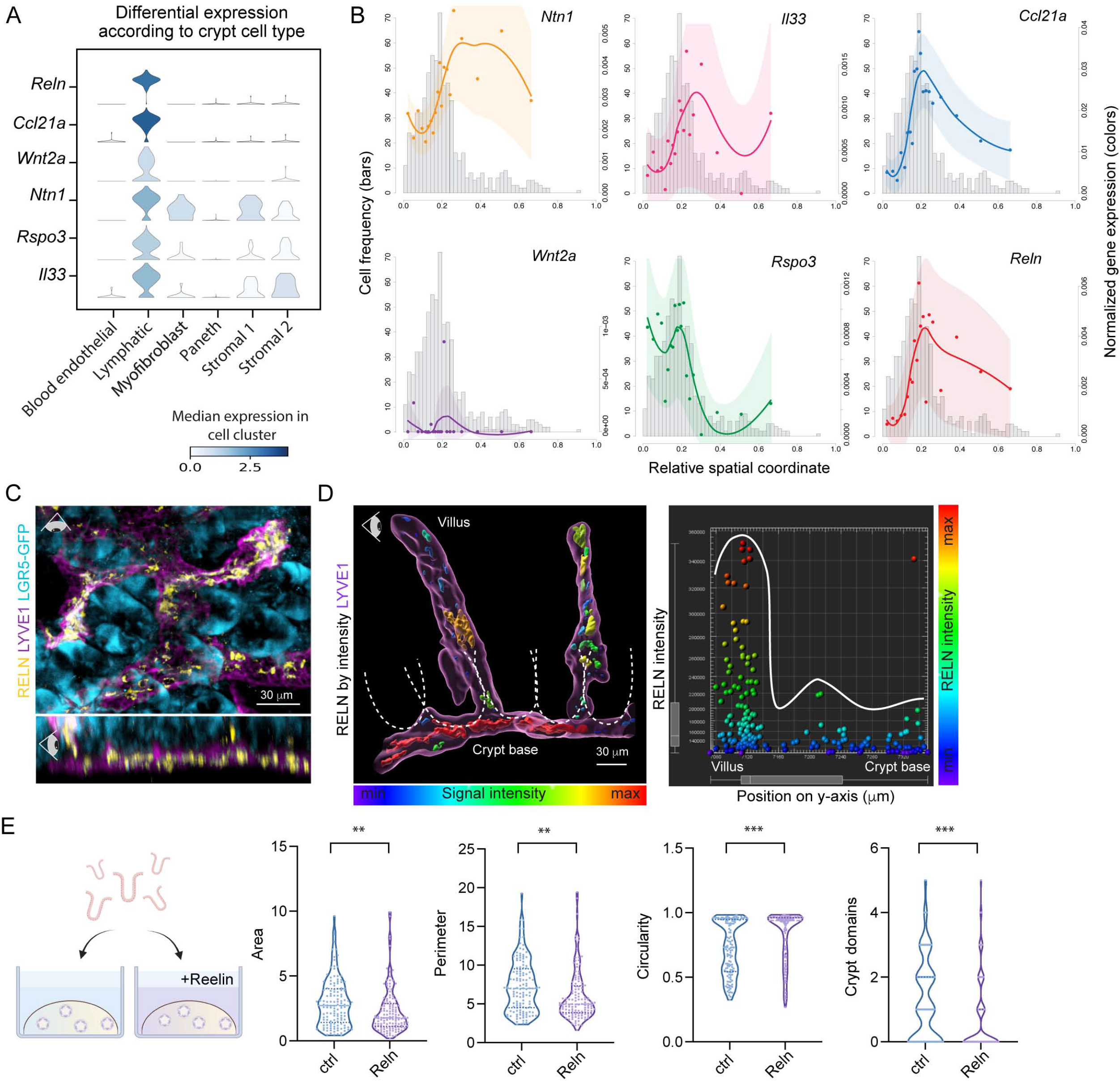
Localized expression of the lymphatic:stem cell interactome identifies lymphatic-derived niche factors. **(A)** Violin plots of the log-normalized expression of candidate secreted factors in selected cell types in the mouse small intestinal scRNA-seq dataset. **(B)** SpaceFold cartographic maps of the expression of candidate lymphatic ligand genes normalized by the total gene expression of lymphatics over the crypt-villus axis. X-axes mark the relative spatial coordinate imputed by SpaceFold cartography. Histograms show the distribution of spots containing lymphatics. Y-axes on the left indicate the frequency of the bars in the histograms. Y-axes on the right indicate the normalized expression value, *U̅* of the indicated gene in lymphatics. Each dot represents the expression values averaged over an interval of spots containing lymphatics binned by the coordinate. The x-coordinate of each spot represents the mean of the SpaceFold coordinates in each bin. Lines mark the mean values fitted using local polynomial regression. Shaded areas represent the mean ± 2 standard error. **(C)** Whole-mount immunofluorescence image of small intestinal crypts harboring LGR5^+^ stem cells (cyan) that are embedded in a network of LYVE1^+^ lymphatic capillaries (purple) expressing REELIN (yellow). The top image shows the tissue from a bird’s eye view, whereas the bottom panel depicts a side view. **(D)** Left panel: 3D-reconstructed immunofluorescence whole-mount image of the lymphatic capillary network (LYVE1^+^, purple) along the crypt-villus axis in the small intestine. Dotted lines outline individual crypts. REELIN expression in lymphatics is color-coded by signal intensity, demonstrating highest intensity at the base of crypts. Right panel: quantification of the signal intensity of REELIN immunofluorescence in the adjacent figure panel plotted against the y-position. Each dot represents a surface spot of REELIN fluorescence, generated in Imaris, color-coded by its’ REELIN signal intensity and plotted along the crypt-villus axis **(E)** Left panel: schematic of organoid experiments with recombinant murine REELIN. Right panel: violin plots of the area, perimeter, circularity and number of crypt domains in organoids cultured in the presence or absence of murine recombinant REELIN for 4-5 days. Data were generated from *n =* 3 independent experiments. *** indicates a p-value of 0.0001 to 0.001, ** indicates a p-value of 0.001 to 0.01 (Unpaired two-tailed Student’s *t*-test).

R-SPONDIN-3, encoded by *Rspo*3, binds to LGR5 and enhances WNT signaling in intestinal stem cells (Mah et al., 2016; Yan et al., 2017). Although it was initially identified as a factor produced by mesenchymal cells (Greicius et al., 2018), *Rspo3* expression was recently reported to be expressed by lymphatics as well (Ogasawara et al., 2018). Our transcriptomic data clearly corroborate crypt-based lymphatics as the major source of this essential ISC ligand (Figure S6A). While WNT2 has not been studied in the context of the crypt lymphatics, it added to the list of canonical WNTs expressed by Paneth and mesenchymal cells of the niche (David et al., 2020; Mah et al., 2016). Given the importance of WNT-Frizzled signaling in ISC maintenance, the prominence of these factors in crypt-based LECs heightened the importance of lymphatics as a critical new component of the ISC niche.

While the roles of R-SPONDINs and WNTs in ISC function were well established, REELIN had not been previously described as an ISC niche factor, but rather as a large extracellular matrix protein involved in neuronal migration (D’Arcangelo et al., 1995) and cardiac remodeling (Liu et al., 2020). Given that REELIN’s primary receptors, *Vldlr/Lrp8* and *Itgb1* (D’Arcangelo et al., 1999; Dulabon et al., 2000), were all expressed by ISCs and *Reln* was selectively expressed by crypt-based lymphatics (Figures S6A-S6B), it surfaced as a good candidate for eliciting the effects of LECs on ISC maintenance and differentiation.

As judged by immunofluorescence, REELIN was expressed in our cultured LECs, and tissue clearing and whole-mount imaging confirmed its enriched expression in the LECs that concentrated along the crypt base capillary network in both small and large intestine (Figures 6C-6D, S6C-S6D and Video S7). Most strikingly, the addition of recombinant REELIN to standard organoid cultures recapitulated the phenotype of organoids cocultured with LECs and/or cultured in LEC-conditioned media, with REELIN-treated organoids demonstrating reduced size, increased circularity and a decreased number of crypt domains (Figure 6E). These data underscored the functional importance and physiological relevance of the lymphangiogenic secretome in governing ISC behavior and maintaining stemness in the intestinal crypt niche. Furthermore our study illuminated the power of integrated transcriptomic and high-resolution spatial analysis in dissecting the complexity of stem cell niches and in identifying novel niche components.

## Discussion

### Lymphatics and the Intestinal Stem Cell Niche

Inverse gradients of WNTs and BMPs as well as other signals are known to be generated by the diverse cellular constituents of the intestinal crypt niche to impact ISC self-renewal and differentiation in health and disease (Beumer and Clevers, 2021). Yet, the comprehensive nature of these cues and how they are dynamically coordinated to orchestrate stem cell behavior across tissues remain incompletely understood. We previously discovered that lymphatic capillaries form a dynamic niche for the hair follicle stem cells in the skin, which undergo cyclical bouts of quiescence and regenerative activity to drive the hair cycle (Gur-Cohen et al., 2019). In contrast to the skin, the focus of the intestinal lymphatic network has been on the lacteal, a central lymphatic capillary enwrapped by a blood vascular network, that extends from the intestinal submucosa into the villus and functions in transporting dietary lipids and nutrients, immunosurveillance and fluid balance regulation. Although connected to small intestinal lacteals, the lymphatic capillary network below the crypts throughout the intestine has been largely ignored. In this regard, the intimate relation between ISC-harboring crypts and lymphatics as revealed by our 3D-imaging and substantiated by ultrastructural analyses was both notable and intriguing.

The lymphatic-organoid coculture system that we established revealed a potent effect of lymphatic endothelial cells on maintaining the self-renewal ability and fitness of ISCs while suppressing their differentiation. We traced this feature back to factors secreted by lymphatics, as LEC-conditioned media was sufficient to elicit these phenotypic changes in organoids. Importantly, when pre-conditioned ISCs were grown in normal organoid culture conditions, they retained superior tissue-regenerating activity compared to ISCs cultured in normal media. Upon removal of LEC-secreted factors in conditioned media, those ISCs were able to progress normally along their differentiation lineages, though.

### Lymphatics as a Signaling Hub for Tissue Stem Cells

Instead of taking a proteomic approach to identify the lymphatic factors involved, we devised a transcriptomic strategy to arrive at candidates which we could then interrogate functionally. BayesPrism integrated our 10X Visium spatial transcriptomic with our scRNA-seq data, in a manner that not only revealed cell type fractions, but also cell type-specific gene expression profiles per spot. Taking advantage of a scRNA-seq based reference atlas to deconvolve low-resolution spatial transcriptomic data with our new SpaceFold approach, we computationally reconstructed the spatial organization of the crypt-villus unit. This analysis unveiled a high-resolution spatial map of each cell within the small and large intestine and their spatially determined gene programs. By enriching for lymphatic endothelial cells and rare LGR5^+^ progenitor cells, these maps became all the more powerful. This was illustrated by our ability to dissect transcriptional heterogeneity within the lymphatic endothelial network across the intestinal crypt-villus axis, and to identify factors produced specifically by lymphatic capillaries at the crypt base.

This strategy worked beautifully in guiding us to the unique lymphangiocrine signals that directly instruct ISC development and maturation. Through localizing niche signals and/or their ISC receptors, a mechanism surfaced, whereby ISC progeny displaced from their niche signals execute a program of differentiation. This was corroborated by our findings *in vitro*, when re-propagating organoids grown in LEC-conditioned media. Our cartographic approach clarified prior ambiguities regarding the major source of *Rspo3,* encoding a key WNT-enhancing niche signal for LGR5^+^ ISCs (Greicius et al., 2018; Ogasawara et al., 2018), and identified REELIN as an additional lymphatic candidate factor signaling to ISCs within the crypt niche. Indeed, in our organoid models, RSPONDINs (RSPO-1 is included in our ENR organoid media) are known to be essential for ISC maintenance and as we showed here, REELIN also had a direct and distinct impact on organoid development and maturation.

In revisiting the HFSC niche, where surrounding lymphatic capillaries play a key role in stem cell maintenance and hair regeneration (Gur-Cohen et al., 2019), it is intriguing that *Reln* (in LECs) and *Lrp8/Vdlr/Itgb1* (in HFSCs) are also expressed, suggesting a conserved role for REELIN signaling in tissue stem cell niches (Gur-Cohen et al., 2019; Kalucka et al., 2020). Additionally, despite differences in stem cell activities and tissue regeneration in the intestine and skin, lymphatics seem to function similarly, namely by governing the maintenance and regenerative activities within the stem cell niche. In this regard, it is interesting to note that despite some conserved features of stem cell niches, as we’ve unearthed here, others appear to be tailored to suit the particular needs of each tissue.

### The Power and Universality of Our Computational Strategy in Unearthing Intercellular Interactions Within Tissues

While beyond the scope of the current study, the high-resolution spatial map we charted here and the associated deep single-cell profiling of small and large intestinal cells now offers a rich dataset to query additional insights into spatially defined cell types and gene expression programs as well as cellular interactions within the intestine. While most previous studies have focused only on inferring cell type fractions from spatial transcriptomics, our study has highlighted the importance of also inferring spatially resolved cell-type specific gene expression. Moreover, this method can be applied to spatial transcriptomic data from other tissues comprised of spatially ordered units as well. In this regard, two features of BayesPrism make it highly generalizable to additional spatial transcriptomic data sets: (1) It does not rely on the cell type proportions observed in the scRNA-seq data, making it particularly effective for datasets with enriched rare cell types and (2) it is robust to technical and even biological variation and therefore does not require perfectly matched samples that were profiled via scRNA-seq and spatial transcriptomics.

Taking advantage of the stereotypical crypt-villus and crypt units, SpaceFold was able to compress our transcriptomic data onto a 1D spatial axis, facilitating our analysis of spatially restricted gene expression in the intestine. SpaceFold can be applied similarly to other stereotypical tissue structures (e.g. hair follicles) and can be extended to project each spot onto two dimensions, to construct a 2D cartography of more complex tissue units (e.g. liver lobules).

### Broader Implications for Lymphatic-ISC Interactions: Looking Ahead

Although we focused here on direct communication between lymphatic endothelial cells and ISCs, lymphatics are also likely to mediate other crypt dynamics, including trafficking of niche-specific signaling molecules, fluid drainage and immune cell crosstalk with ISCs. Indeed, while immune cells can directly signal to stem cells to orchestrate pathogen responses (Biton et al., 2018; von Moltke et al., 2016), immune-mediated damage of ISCs is a common feature of graft-versus-host disease, in which lymphoid organs are damaged (Fu et al., 2019), suggesting an important role for immune cell drainage from the niche via lymphatics.

With the diversity of roles that lymphatics can play, their association with ISCs is also likely to have important consequences in disease states. In this regard, inflammatory diseases of the intestine have been associated with lymphatic vessel dilation, submucosal edema, lymphatic obstruction, lymphadenopathy, lymphangiectasia, and lymphangiogenesis (Alexander et al., 2010; D’Alessio et al., 2014; Jang et al., 2013; Van Kruiningen and Colombel, 2008). Additionally, obesity alters LEC density, proliferation, and permeability (Zhang et al., 2018). Any or all of these perturbations in lymphatics could impact ISCs directly. To guide therapies in the future, it will be important to interrogate how the lymphatic transcriptome is altered in disease states, how it interprets inflammatory and metabolic cues, and how it relays them to the ISC niche.

## Acknowledgments

We thank the Rockefeller University’s Resource Centers: Flow Cytometry (S. Mazel, director), Bioinformatics (T. Carroll, W. Wang); Genomics (C. Zhao, director) and Memorial Sloan Kettering Cancer Center’s Integrated Genomics Operation (N. Mohibullah) and Molecular Cytology Facility (K. Manova-Todorova, Y. Romin, N. Fan). We thank R. Longman for human biopsies and helpful discussions. We thank R. Chaligné, D. Mucida, S. Josefowicz, S. Tavazoie, A.Hanash, D. Artis and R.P. Kataru for helpful discussions. We thank E.F. laboratory members for: technical assistance (E. Wong, M. Nikolova, J. Racelis, P. Nasseir); administrative assistance (C. Long); mouse handling and experiments (M. Sribour, L. Hidalgo, J. Levorse, L. Polak); helpful discussions (N. Gomez, S. Ellis, M. Parigi, A. Gola, M.D. Abdusselamoglu, S. Baksh, S. Liu, M. Tierney, J. Novak, C. Xu). We thank J. Shin in the Mehrara lab for technical assistance. R.E.N. is a Burroughs Wellcome CAMS recipient and a clinical scholar at The Rockefeller University and receives support in part by the National Center for Advancing Translational Sciences, NIH, through The Rockefeller University (grants UL1 TR001866 and KL2TR001865). S. G. C. is a Robin Chemers Neustein fellow (2021) and former Human Frontier Science Program (HFSP) LT001519/2017 and European Molecular Biology Organization (EMBO) ALTF 1239-2016 fellow. T.C. is a Croucher Foundation postdoctoral fellow and Damon Runyon Quantitative Biology postdoctoral fellow. M.S. is a Rockefeller University Women & Science (2020) and Boehringer Ingelheim Fonds (2021) graduate fellow. E.F. is an Investigator with HHMI. The work was supported by grants from the National Institutes of Health (NIAMS R01-AR050452, E.F.) and the STARR Foundation (2019-009, E.F. and B.J.M.).

## Author contributions

Conceptualization E.F., D.P., B.J.M., R.E.N., T.C., S.G-C., M.S.; Methodology E.F., D.P., R.E.N, T.C., S.G-C, M.S.; Computational algorithms D.P., T.C., M.S.; Validation R.E.N., T.C., S.G-C., M.S.; Formal analysis R.E.N., T.C., S.G-C., M.S, H.A.P.; Investigation R.E.N., T.C., S.G-C., L.H., M.S., H.A.P., R.P.K. Resources E.F., D.P., B.J.M., R.E.N; Data Curation R.E.N., T.C., S.G-C., M.S.; Manuscript writing E.F., D.P., B.J.M., R.E.N, T.C., S.G-C., M.S.; Visualization E.F., D.P., R.E.N., T.C., S.G-C., M.S.. H.A.P.; Supervision E.F., D.P., R.E.N., S.G-C.; Project Administration E.F., D.P., R.E.N., S.G-C., M.S.

## Declaration of interests

The authors declare no competing interests. E.F. is on the scientific advisory boards of L’Oréal and Arsenal Biosciences. D.P. is on the scientific advisory board of Insitro. B.J.M is an advisor to PureTech Corp and recipient of a investigator-initiated research award from PureTech Corp and recipient of an investigator-initiated research award from Regeneron.

## Supplemental figure captions

**Figure S1.**
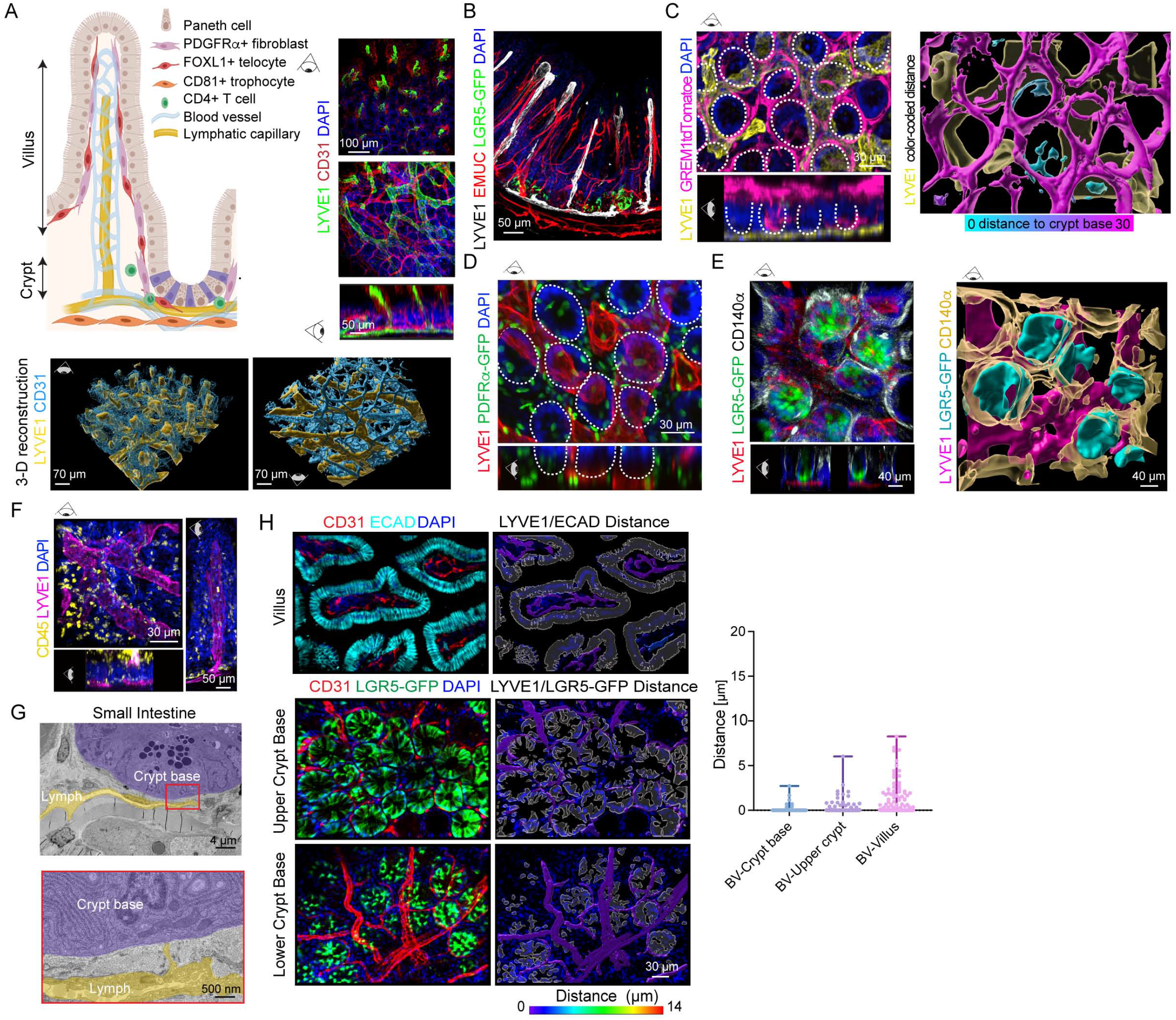
Lymphatics integrate with neighboring intestinal stem cell niche cells to access the crypt base. **(A)** Left: schematic of the intestinal crypt niche, indicating cell types approximating the crypt base and their integration with the lymphatic niche. Right: 3D images of the small intestinal vasculature in the villus illustrating LYVE1^+^ lymphatics (green) and CD31^+^ blood endothelial cells (red). Bottom panel: representative 3D-volumetric reconstruction of vascular structures including lymphatics (LYVE1^+^, yellow) and blood vessels (CD31^+^, blue). **(B)** 3D image of the lymphatic vasculature (LYVE1^+^, white), blood vessels (EMUC^+^, red) and crypt-based LGR5^+^ stem cells (green) in the small intestine of Lgr5-eGFP-IRES-CreERT2 mice from a side view. **(C)** 3D images of GREM1^+^ stromal cells, lymphatics (LYVE1^+^) and crypts (DAPI^+^, outlined by dotted lines) in Grem1-CreERT2-tdTomato^flox/flox^ mice. The left panel shows the original immunofluorescent image, while the right panel shows 3D-reconstructed GREM1^+^ stromal cells color-coded by their distance to the crypt base. **(D)** 3D image of small intestinal crypts demonstrating the relationship between PDGFR*α*/CD140a^+^ stromal cells, the lymphatic vasculature (LYVE1^+^) and DAPI^+^ crypts (white dotted outline) in PDGFR*α*-H2B-eGFP mice, in which nuclei of PDGFR*α*^+^ cells are shown in green. **(E)** 3D image of small intestinal tissue stained for PDGFR*α*/CD140a, LYVE1, and LGR5-GFP in Lgr5-eGFP-IRES-CreERT2 mice demonstrating the relationship between PDGFR*α*^+^ mesenchymal niche cells, lymphatics and crypt-based stem cells. 3D reconstruction is in the right side panel. **(F)** 3D images of mouse small intestinal tissue demonstrating the spatial relationship between CD45^+^ immune cells, LYVE1^+^ lymphatics and epithelial cells (DAPI^+^) of intestinal crypts and villi with the left panel illustrating the crypt region and the right panel illustrating the full crypt-villus axis and a lymphatic lacteal. **(G)** EM image demonstrating the crypt base (purple, identified by granules within Paneth cells) and lymphatic capillaries (yellow). The inset (red box) demonstrates a lymphatic endothelial cell projection towards the crypt base. **(H)** 3D images of 5 um z-stacks of the mouse small intestinal blood vasculature and epithelial cells (villus) or intestinal stem cells (upper and lower crypt base). Quantification of the distance between vascular surfaces and the epithelium was done in each of the three regions. (*n=* 9 mice with 3 images/region/mouse). Ns means non-significant (one-way ANOVA with Tukey’s multiple comparisons). Figures A-G are representative of *n =* 3 mice/group with ≥ 3 images/mouse/region imaged.

**Figure S2.**
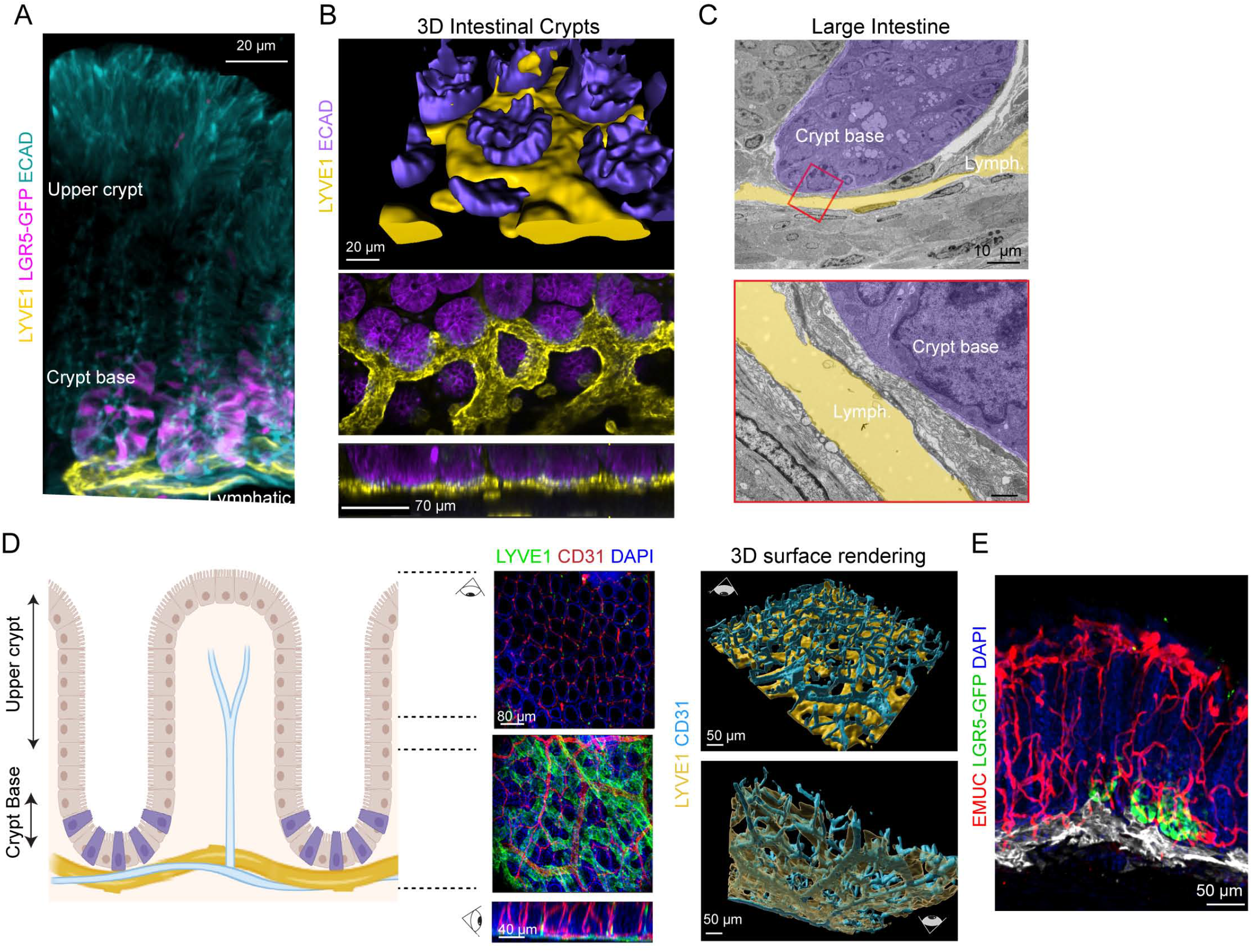
Large intestinal lymphatics are tightly associated with the crypt base. **(A)** 3D image demonstrating the association between crypt-based intestinal stem cells (LGR5-GFP^+^) and lymphatic vasculature (LYVE1^+^) in the mouse large intestine. **(B)** 3D rendering of large intestinal crypts (ECAD^+^) and lymphatics (LYVE1^+^) in the mouse large intestine. **(C)** EM images demonstrating the crypt base (purple) and lymphatic capillaries (Lymph., yellow) in the mouse large intestine. **(D)** 3D images of the large intestinal vasculature in the upper crypt and crypt base in the large intestine with a schematic diagram on the left demonstrating LYVE1^+^ lymphatics and CD31^+^ endothelial cells. Right side panels are with representative 3D volumetric reconstruction of vascular structures. **(E)** 3D image of the lymphatic vasculature (LYVE1^+^), blood vessels (EMUC^+^) and crypt-based LGR5^+^ stem cells in the large intestine of Lgr5-eGFP-IRES-CreERT2 mice. Images are representative of *n =* 3 mice/group with ≥ 3 images/mouse/region imaged.

**Figure S3.**
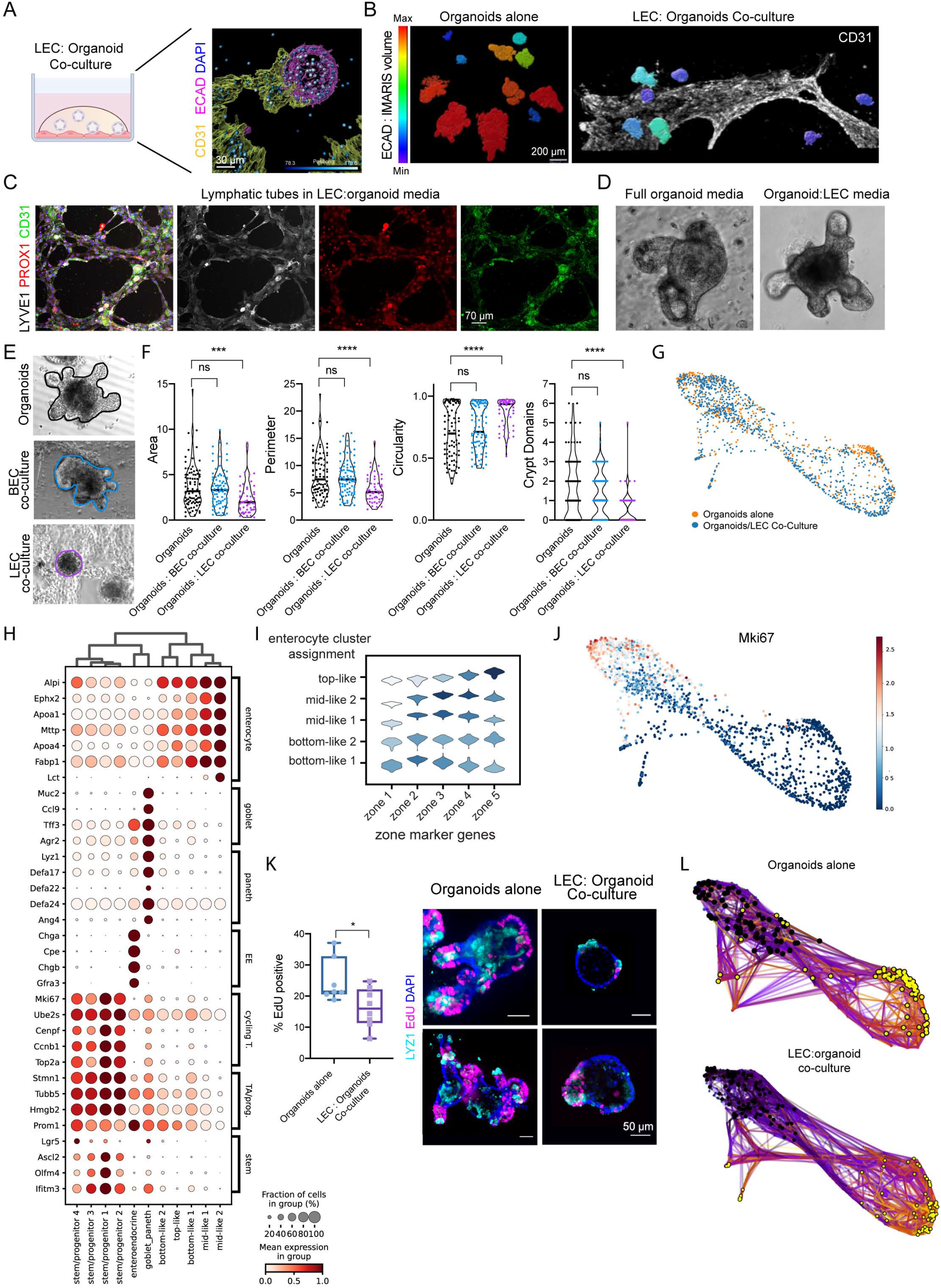
Phenotypic characterization of the intestinal organoid:lymphatic endothelial cell coculture system. **(A)** Scheme of the LEC:organoid coculture system with a 3D-reconstructed immunofluorescence image of CD31^+^ LECs (yellow) interacting with ECAD^+^ small intestinal organoids (purple) in culture. Nuclei are color-coded by relative z-position. **(B)** Representative images of small intestinal organoids grown in the absence (left panel) or presence (right panel) of CD31^+^ LECs (white). 3D-reconstructed surfaces of organoids are color-coded according to their volume. **(C)** Representative immunofluorescence images of cultured LECs at passage 5. LECs maintain common endothelial markers such as CD31 (green), LYVE1 (white) and PROX1 (red) while grown in 50:50 co-culture media. **(D)** Representative brightfield images of small intestinal organoids cultured in 100% organoid medium (left) and 50%:50% LEC:organoid medium (right). **(E)** Representative images of small intestinal organoids cultured alone or cocultured with LECs and BECs. Tracing of organoid shape was done in ImageJ for subsequent quantification of morphological parameters in F. **(F)** Violin plots of the area, perimeter, circularity and number of crypt domains of small intestinal organoids cultured alone, cocultured with BECs or LECs (*n =* 3 independent experiments). **** indicates a p-value of <0.0001, *** indicates a p-value of 0.0001 to 0.001 and ns is non-significant (one-way ANOVA with Tukey’s multiple comparisons). **(G)** Force-directed layout of scRNA-seq data of small intestinal organoids that were cultured in the absence (orange dots) and presence (blue dots) of LECs. **(H)** Dot plot of marker genes used for cell type-specific cluster assignment of sequenced epithelial cells from organoids cultured in the presence or absence of LECs. The size of the dots represents the fraction of cells with non-zero expression in each group, while the color indicates the log-transformed normalized UMIs for each gene standardized to between 0 and 1. Final cluster IDs are listed at the bottom, genes are indicated on the left, while their general classification according to cell type is indicated on the right. EE = enteroendocrine cell, cycling T. = cycling transit-amplifying (TA) cell, TA/prog. = TA/progenitor cell **(I)** Violin plot of enterocyte zone signatures (x-axis) in absorptive enterocyte clusters (y-axis). **(J)** *Mki67* gene expression projected onto the UMAP plot, marking cycling cells. Cells are colored by the log-transformed normalized UMIs of *Mki67*. **(K)** Boxplot showing the percentage of 5-Ethynyl-2′-deoxyuridine (EdU^+^, magenta) cells after a 10 minute-pulse in small intestinal organoids cultured in the absence or presence of LECs. (*n =* 2 independent experiments). * indicates a p-value of <0.05 (Unpaired two-tailed Student’s t-test). Representative images of quantified organoids are on the right (LYZ1^+^ Paneth cells in light blue, DAPI^+^ nuclei in blue and EdU^+^ cells in magenta). **(L)** Terminal cell states predicted by a random walk simulation in each condition. Cells were visualized using the same force-directed layout as in G. Black dots represent the cells sampled as the starting points, which were chosen from the cells labeled as the stem/TA cluster in each condition. Yellow dots represent cells of the end points. Each line represents a simulated random walk between the cells connected by it.

**Figure S4.**
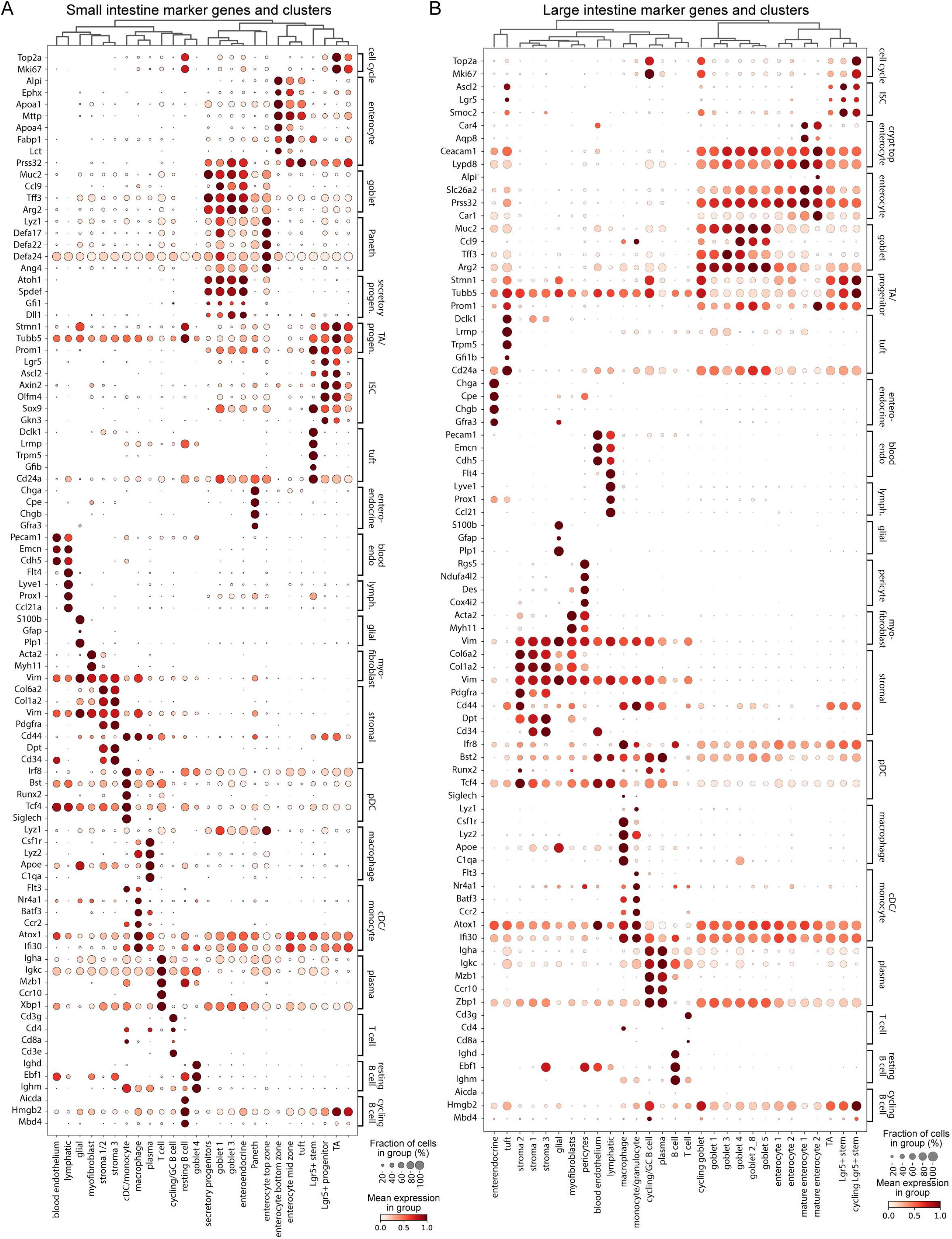
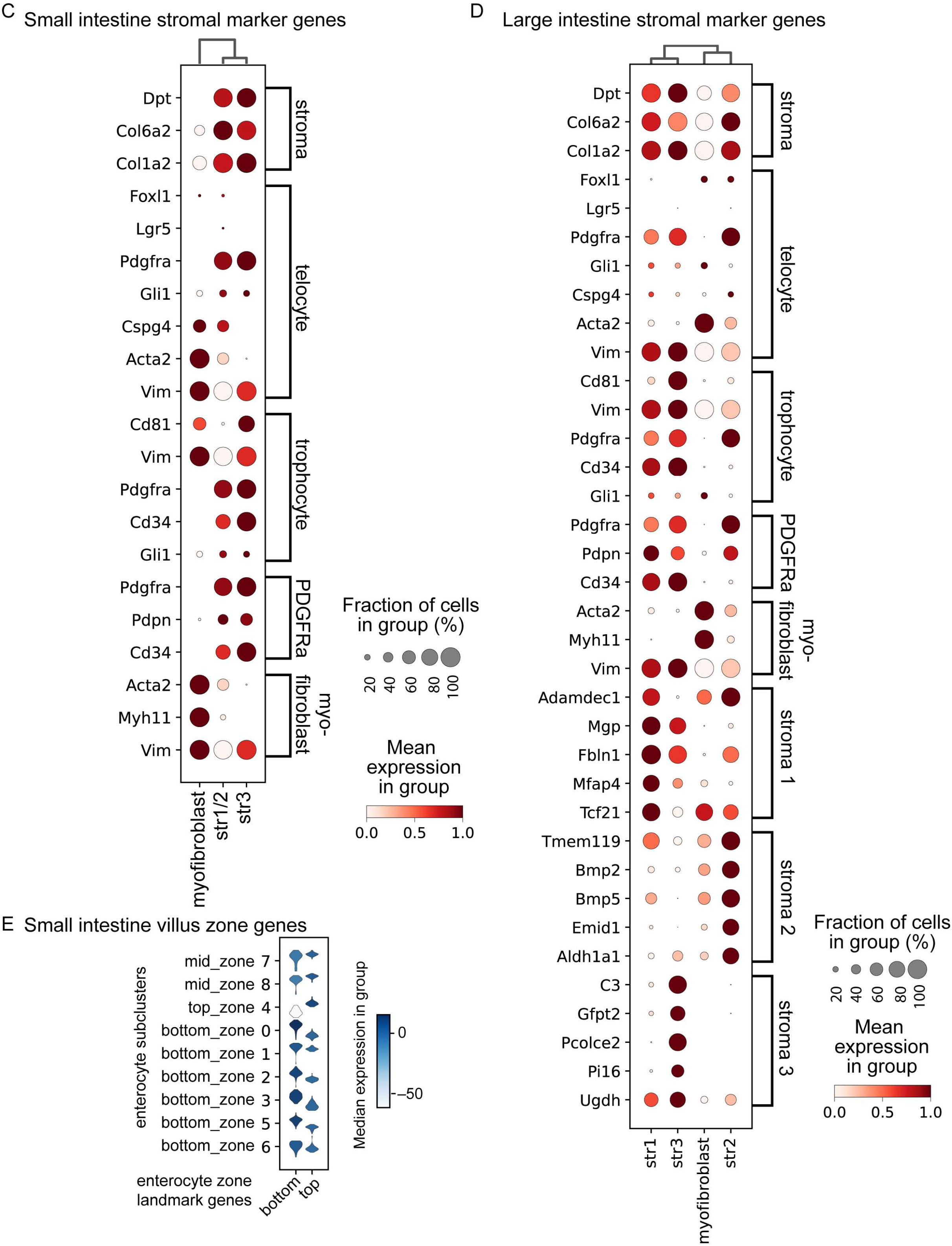
Dot plots show the expression of marker genes in each cluster of cells in the *in vivo* sequencing datasets. **(A-B)** Expression of marker genes in all final clusters from the murine small **(A)** and large intestinal tissue dataset **(B)**. The size of the dots represents the fraction of cells with non-zero expression in each group, while the color indicates the log-transformed normalized UMIs for each gene standardized to between 0 and 1. **(C-D)** Dot plots showing the expression of marker genes for small (C) and large intestinal (D) stromal niche components as described in the literature. For all the plots in A-D final cluster IDs are listed at the bottom, genes are indicated on the left and their classification according to cell type is shown on the right. Lymph = lymphatics, pDC = plasmacytoid dendritic cell, cDC = classical dendritic cell, GC B cell = germinal center B cell, TA = transit-amplifying cell. **(E)** Distribution of the expression of the mean z-scores of the top and bottom landmark genes reported by Moor et al. in each enterocyte subcluster of the small intestine dataset.

**Figure S5.**
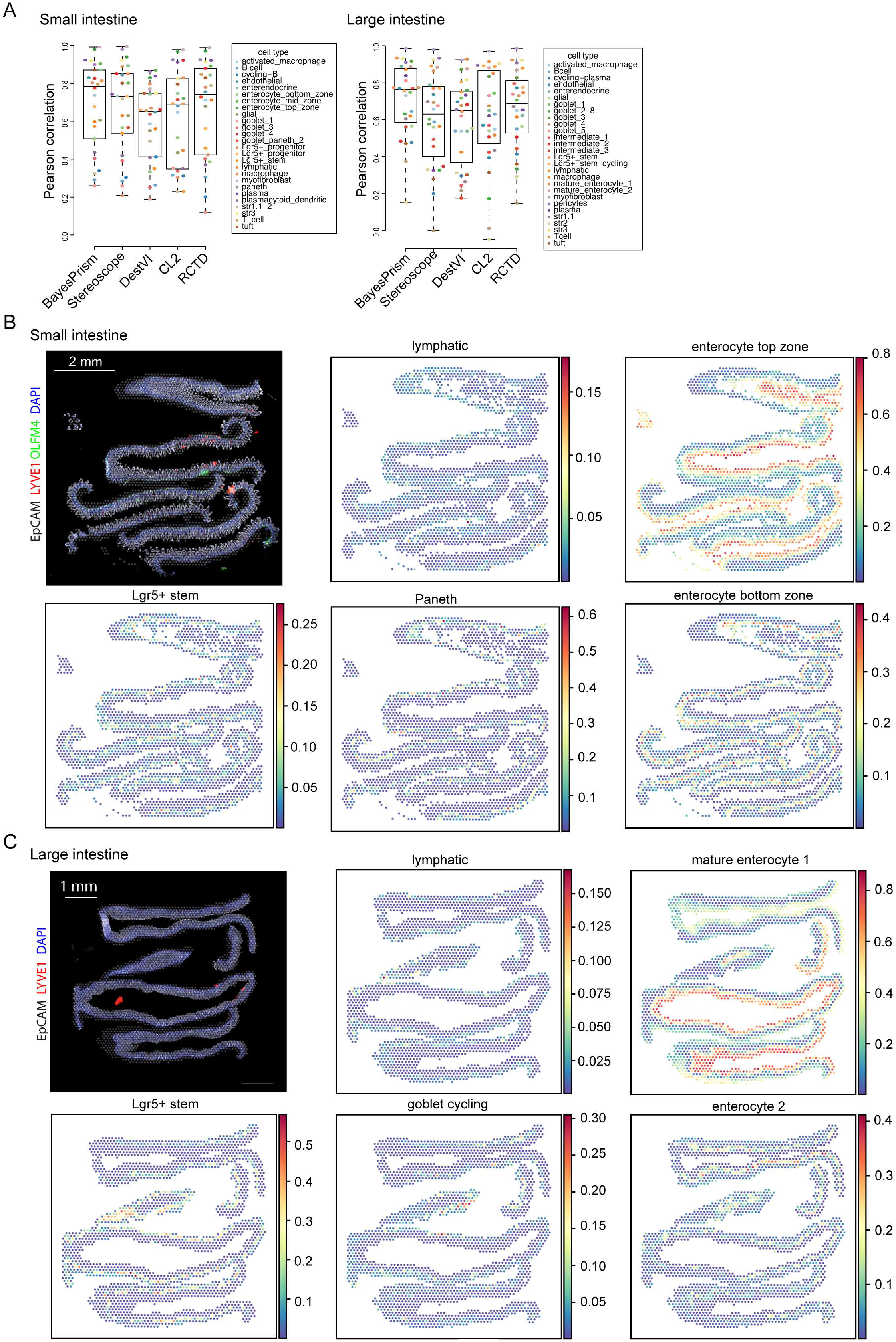
Benchmarking BayesPrism with other tools for deconvolving spatial transcriptomic data. **(A)** Boxplots show Pearson’s correlation between cell type fractions inferred by each method using the same set of cell type marker genes, and the fraction of reads from the top 50 differentially expressed genes in each cell type. Boxes mark the 25th percentile (bottom of box), median (central bar), and 75th percentile (top of box). Whiskers represent extreme values within 1.5-fold of the interquartile range. Dots are colored according to their cluster assignment/cell type ID. **(B-C)** Immunofluorescence image of the mouse small intestinal (the top-left panel in B) and large intestinal (the top-left panel in C) tissue section used for spatial transcriptomic (10X Visium) profiling. EpCAM*^+^* epithelial cells are in white, LYVE1*^+^* lymphatic in red and OLFM4^+^ small intestinal stem cells in green. The overlaid gray dots on immunofluorescent images represent the 50 µm diameter RNA-capture areas. The rest of the panels show the fraction of representative cell types with known spatial distributions inferred by BayesPrism (2nd to 5th panel in B and C).

**Figure S6.**
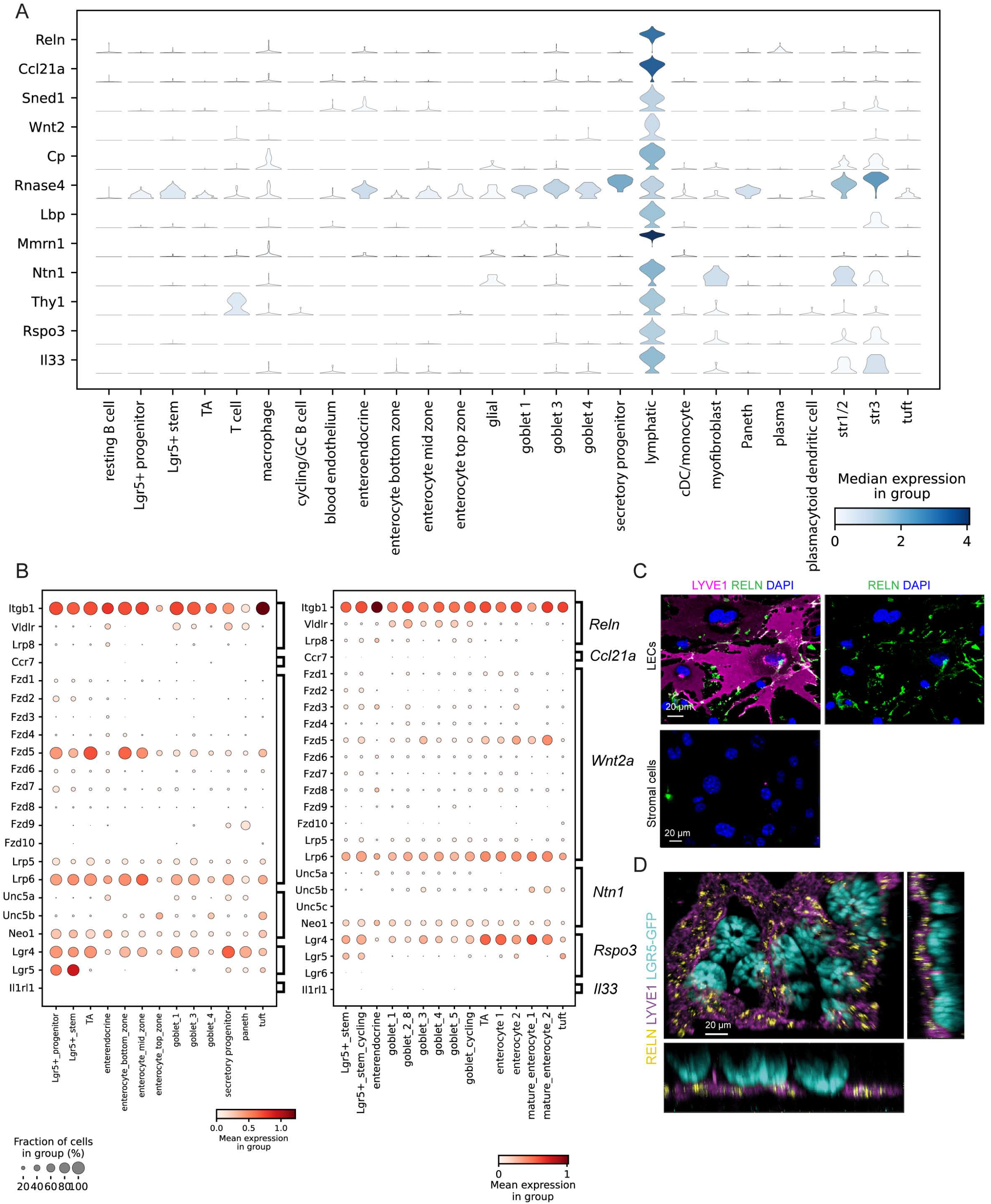
Expression of lymphangiocrine factors across the intestine. **(A)** Distribution of log-normalized UMIs of candidate lymphangiocrine factors among small intestinal cell types computed using the small intestinal scRNA-seq data shown in figure 4B. **(B)** Expression of putative receptors for selected lymphangiocrine factors (indicated on the right) among small (left) and large (right) intestinal cell types computed using the small and large intestinal scRNA-seq data shown in figure 4B and C. The size of dots represents the fraction of cells with non-zero expression in each group, while the color indicates the log-transformed normalized UMIs for each gene standardized to between 0 and 1. Receptor genes are indicated on the left y-axis, cluster IDs are listed at the bottom. **(C)** Immunofluorescence of REELIN staining in cultured lymphatic endothelial cells and stromal cells, which do not produce REELIN (negative control). **(D)** Whole-mount immunofluorescence image of mouse large intestinal tissue showing LGR5^+^ stem cell-harboring crypts (cyan) sitting atop a network of LYVE1^+^ lymphatic capillaries (purple), which express RELN (yellow).

## Multimedia files

**Video S1. The vascular architecture of the small intestine.** Related to Figure 1 and Figure S1. Whole-mount imaging and 3D reconstruction of the small intestinal vasculature in the villus illustrating LYVE1+ lymphatics (green) and CD31+ blood endothelial cells (red). Volumetric reconstruction of lymphatics (LYVE1+) in yellow and blood vessels (CD31+) in light blue.

**Video S2. Lymphatic capillaries are highly associated with the crypt-based stem cells of the small intestine.** Related to Figure 1. Whole mount imaging and 3D reconstruction of small intestinal LYVE1+ lymphatic capillaries (red) nesting LGR5+ crypt-based stem cells (green), and volumetric reconstruction of lymphatics (LYVE1+, purple) and stem cells (LGR5+, yellow).

**Video S3. Lymphatic capillaries are highly associated with crypts in the large intestine.** Related to Figure S2. Whole-mount imaging and 3D reconstruction of large intestinal LYVE1+ lymphatic capillaries (white) and CD31+ blood endothelial cells (red) around crypt-based epithelial cells (ECAD+, green). Volumetric reconstruction indicating lymphatics (LYVE1+, yellow) and crypt-based epithelial cells (purple).

**Video S4. Lymphatic capillaries are nesting crypt-based stem cells of the human colon.** Related to Figure 1. Whole-mount imaging and 3D reconstruction of a human intestinal biopsy, demonstrating association between lymphatics (LYVE1+, red) and epithelial cells of the intestinal crypt base (ECAD+, green) in the distal colon. Volumetric reconstruction indicates lymphatics (LYVE1+, yellow) and crypt-based epithelial cells (purple).

**Video S5: Interaction of an intestinal organoid with LECs in coculture conditions.** Related to Figure 2 and Figure S3. Representative z-stack intervals of organoids (EpCAM+, green) grown on top of lymphatic endothelial cells (CD31+, white), cultured in 50% LEC:50% organoid medium.

**Video S6: Intestinal organoid associated with LECs in coculture conditions. Related to** Figure 2 **and Figure S3.** Confocal 3D imaging and volumetric reconstruction of intestinal organoids (ECAD+, Green; reconstructed IMARIS volume in magenta) associated with lymphatic endothelial cells (CD31+, red; reconstructed IMARIS volume in yellow), cultured in 50% LEC:50% organoid medium.

**Video S7: Crypt-based lymphatic capillaries highly express REELIN.** Related to Figure 6 and Figure S6. Whole-mount imaging and 3D reconstruction of the mouse large intestine showing LGR5+ stem cell-harboring crypts (cyan) sitting atop a network of LYVE1+ lymphatic capillaries (magenta), which highly express RELN (yellow).

## Methods

### RESOURCE AVAILABILITY

#### Lead contact

Further information and requests for resources and reagents should be directed to and will be fulfilled by the lead contact, Dr. Elaine Fuchs.

#### Materials availability

Will be provided upon request and available upon publication.

#### Data and code availability

All data that support the findings of this study are available within the paper and its supplementary files. All single-cell and spatial transcriptomic data generated within this study have been deposited in the Gene Expression Omnibus (GEO) repository with the accession code GSE190037. All codes will be uploaded on the Pe’er lab github website (https://github.com/dpeerlab/SpaceFold_paper).

### EXPERIMENTAL MODEL AND SUBJECT DETAILS

#### Animals

C57BL/6 mice, Lgr5-eGFP-IRES-CreERT2 (B6.129P2-*Lgr5^tm1(cre/ERT2)Cle^*/J), PDGFR*α*-H2B-eGFP (B6.129S4-*Pdgfrα^tm11(EGFP)Sor^*/J)and ROSA^mT/mG^ (*Gt(ROSA)26Sor^tm4(ACTB-tdTomato,-EGFP)Luo^*/J)) mice were purchased from the Jackson Laboratories. Both male and female mice were used. Mice were maintained in the Association for Assessment and Accreditation of Laboratory Animal Care-accredited animal facility of The Rockefeller University (RU), and procedures were performed with Institutional Animal Care and Use Committee (IACUC)-approved protocols. Mice of all strains were housed in an environment with controlled temperature and humidity, on a 12-hour light cycle, and fed regular rodent’s chow. The numbers of animals used for each experiment is presented in the figure legends legends as *n =* x mice per group, per time point analyzed. Animals were used at 6-12 weeks of age for experiments unless indicated otherwise. (B6.Cg-Tg(Grem1-cre/ERT)3Tcw/J) mice were a gift from Dr. Timothy C. Wang (Columbia University, New York). ROSA^mT/mG^ (*Gt(ROSA)26Sor^tm4(ACTB-tdTomato,-EGFP)Luo^*/J) mice were used for the majority of crypt isolation (organoid) experiments.

#### Human tissues

Deidentified intestinal biopsy specimens were obtained from the Jill Roberts Inflammatory Bowel Disease Center at Weill Cornell Medical College, Department of Gastroenterology (New York, NY) in compliance with federal and state laws, and National Institute of Health guidelines.

#### Primary Cell Culture

##### Mouse small intestinal organoid culture

Organoids were generated from isolated crypts of the mouse small intestine as described with some alterations (O’Rourke et al., 2016). Briefly, three-quarters of the small intestine was flushed with ice-cold phosphate-buffered saline (PBS), opened longitudinally and scraped twice with a glass slide to remove villus epithelium. The tissue was cut into ∼ 5-10 mm pieces, which were washed 5-6 times in ice-cold PBS with 1.5 mM DTT (Sigma). Remnant villi were removed by shaking tissue pieces at 4°C in an initial 5 minute incubation PBS with 10 mM RNase-free EDTA pH 8.0 (Invitrogen). Crypts were collected and filtered through 70 µm cell strainers in subsequent fractions, for which the pieces were incubated in PBS + 5 mM EDTA for 10 minutes shaking at 4°C. The best fractions were pooled and centrifuged at 4°C in subsequent spins at 290 g and 200 g for 6 minutes each. 300-500 crypts were plated per dome unless noted otherwise.

Organoids were maintained in Advanced DMEM/F12 medium with L-glutamine (2 mM), Penicillin-Streptomycin (100 µg/ml), HEPES buffer (10 mM) and N-Acetylcysteine (1 mM). Growth factors facilitating organoid growth were added and included recombinant murine EGF (50 ng/ml), recombinant murine Noggin (50 ng/ml), recombinant human R-Spondin-1 (500 ng/ml) (ENR medium), unless otherwise specified. To avoid fungal contamination Normocin (InvivoGen) antimicrobial reagent was added at 1:200 to ENR organoid medium. Indicated numbers of crypts or single cells were plated in growth-factor-reduced Matrigel (Corning, R&D) mixed 1:1 with ENR medium. The media was replaced every 2 days and for the first two days in culture after a crypt isolation ROCK inhibitor (1:5000) was added to ENR medium.

To passage organoids, the media was aspirated and replaced with Gentle Cell Dissociation Reagent (Stemcell Technologies) on days 6-7. Matrigel domes were collected in a precoated 15 ml falcon tube and incubated in Gentle Cell Dissociation Reagent for 10 minutes on a rocker at room temperature. Organoids were spun down in subsequent centrifugation steps at 290 g, 200 g and 100 g for 6 minutes and resuspended in ENR organoid medium mixed 1:1 with Matrigel to plate new domes.

#### Mouse lymphatic endothelial cell culture

Primary mouse lymphatic endothelial cells (LECs) were purchased from Cell Biologics (C57BL/6 Mouse primary dermal lymphatic endothelial cells). Lymphatic endothelial cells were expanded on 100 mm dishes, pre-coated with gelatin (Sigma), in Complete Mouse Endothelial Cell Medium (Cell Biologics). LECs were utilized at passages 4-5 in all experiments.

#### Mouse blood endothelial cell culture

Blood endothelial cells (BECs) were derived by sorting alive, CD31^+^, LYVE1^-^, PDPN^-^ single cells from primary mouse endothelial cells (C57BL/6 mouse primary dermal microvascular endothelial cells) purchased from Cell Biologics after *in vitro* expansion. Sorted BECs were used at passages 4-5 for experiments.

### METHOD DETAILS

#### In vitro cell culture

##### Lymphatic endothelial cell-intestinal organoid coculture

For LEC:intestinal organoid coculture experiments, 70,000-100,000 LECs/well were plated on Matrigel-coated 48-well-plates or Millicell EZ 8-well glass slides (Millipore, LabTek). LECs were allowed to adhere and form tubes for 4-6 hours before medium was aspirated and small intestinal crypts or single cells from organoids were plated atop in 20µl Matrigel domes mixed with organoid media.

LEC:intestinal organoid cocultures were maintained in a medium containing 50% Complete Mouse Endothelial Cell Medium and 50% ENR small intestinal organoid medium (50:50 medium). Cocultures initiated from intestinal crypts were maintained for up to 5 days. Cocultures initiated from single intestinal epithelial cells were maintained for up to 10 days.

For experiments with REELIN, intestinal organoids were grown in ENR organoid medium supplemented with 10 nM mouse recombinant Reelin (R&D) daily. If indicated, purified NA/LE Hamster anti-rat CD29 (BD Biosciences) was added at 10 µg/ml for 1 hour prior to adding REELIN.

#### Generation of LEC-and BEC-conditioned media

For experiments with conditioned media, LECs or BECs at 90% confluence were washed 3x in PBS before replacing with 7 ml Complete Mouse Endothelial Cell Medium per 100 mm dish of cells. The medium was collected after 16 or 24 hours as indicated and filtered through a 0.22 µm MBS/Polypropylene vacuum filter unit (MilliporeSigma^TM^ Steriflip^TM^). For experiments a mixture of 50% ENR small intestinal organoid medium and 50% LEC-or BEC-conditioned media was used.

#### Organoid replate assay

Organoids grown in 50:50 medium with LEC-conditioned medium on day 4 or 5 were collected in Gentle Cell Dissociation Reagent as described earlier. To get a single-cell-suspension crypt pellets were washed twice in PBS, resuspended in 200 µl TrypLE Express (Gibco) with DNase I (0.1 mg/ml) and incubated at 37°C for up to 20 minutes with intermittent mixing every 5 minutes. Once the majority of the suspension consisted of single cells PBS with 2% fetal bovine serum (FBS) was added 1:1 and the cells were centrifuged at 1400 rpm for 5 minutes. Cells were counted and plated at 50-100 x 10^3^ cells/dome. Cells were grown in ENR organoid medium with ROCK inhibitor (1:5000) for up to 10 days.

For the replate assay of organoids cocultured with LECs, crypts were originally derived from ROSA^mT/mG^ mice to facilitate sorting of epithelial cells later on. On day 4 or 5 of LEC:organoid coculture cells were collected as described. Pan-epithelial cells (alive, EpCAM^+^, mTomato^+^ single cells) were sorted and plated at 50-100 x 10^3^ cells/dome and cultured as indicated above.

#### Immunofluorescence staining

##### Primary cell immunofluorescence staining

Primary murine lymphatic endothelial cells (Cell Biologics) at passages 4-5 were cultured on Millicell EZ 8-well glass slides (Millipore) pre-coated with gelatin (Sigma) in Complete Mouse Endothelial Medium (Cell Biologics). At 80-90% confluence cells were washed with PBS and fixed in 4% paraformaldehyde (PFA) for 30 minutes at room temperature (RT). Cells were washed twice in PBS and incubated in blocking solution (PBS with 0.3% Triton-X-100 (PBST), 2.5% normal donkey serum, 1% gelatin and 1% bovine serum albumin (BSA)) for 1 hour at RT. Primary antibodies were added in blocking solution for 1 hour at RT. Cells were washed 3 times in PBST, secondary antibodies (1:500) were added in blocking buffer at RT for 2 hours. Cells were washed 3 times in PBST and mounted in 4′,6-diamidino-2-phenylindole (DAPI)-containing ProLong Anti-fade Diamond mountant (Thermo Fisher Scientific).

#### Lymphatic endothelial cell:organoid coculture immunofluorescence staining

For stainings of LEC:organoid cocultures in Matrigel cocultures were washed once in PBS, fixed in 2% or 4% PFA for 30-60 minutes at RT on day 4 or 5, washed 3 times in PBS and permeabilized in PBS with 0.3% Triton-X-100 (PBST) for at least 15 minutes. Cells were blocked for 1 hour at RT in PBST with normal donkey serum (NDS, 2.5%), gelatin (1%) and bovine serum albumin (BSA) (1%) before adding primary antibodies in blocking buffer overnight at 4°C. On the next day the samples were allowed to reach RT for 1 hour before washing 3 times in PBST with 2% NDS and incubating with secondary antibodies (1:200) at RT for 2 hours. Samples were washed at least 3 times in PBST with 2 µg/ml DAPI (1:500). LEC:organoid cocultured that were grown on 8-well glass slides were subsequently mounted with DAPI-containing mountant and a glass coverslip. Cocultures fixed in 48-well-plates were imaged as is in PBS.

Stainings of organoids out of Matrigel were performed as previously described. Briefly, organoids were washed once in PBS before adding 300 µl of ice-cold cell recovery solution (Corning) per well. Organoids were incubated for 30-60 minutes at 4°C until Matrigel was completely dissolved. Well contents were gently resuspended 5-10 times, transferred a 15 ml falcon tube and centrifuged in a volume of 10 ml for 3 minutes at 70 g and 4°C. The pellet was resuspended in 1 ml of ice-cold PFA (4%) and organoids were fixed at 4°C for 45 min. 1 ml of ice-cold PBS with 0.1% Tween-20 was added, organoids were mixed, incubated for 10 minutes and spun down at 70 g. The pellet was resuspended in organoid wash buffer (OWB; PBS + 0.1% Triton-X-100 + 10% SDS + 0.2% BSA) and transferred to a 24-well plate for subsequent steps. Organoids were incubated on a rocker (40 rpm) at 4°C for at least 15 minutes before primary antibodies were added overnight at 4°C. On the next day organoids were washed in OWB in 3 x 2 hour steps. Secondary antibodies (1:500) were added in OWB with 1 µg/ml DAPI (1:1000) and organoids were incubated overnight at 4°C. Washing was repeated on the following day before organoids were transferred to a tube, spun at 70 g and gently resuspended in FUnGI mountant as described by van Ineveld *et al.* Organoids were mounted onto 35 mm glass-bottom dishes (Cellvis) and imaged via confocal microscopy.

#### Intestinal tissue section immunofluorescence staining

For immunofluorescence analysis, the mouse intestine was dissected, flushed with ice-cold PBS and fixed with 4% PFA in PBS for 1 - 2 hours at 4°C, washed three times with PBS and incubated with 30% sucrose overnight at 4°C. The tissue was embedded in OCT (Tissue Tek) the following day. Frozen tissue blocks were sectioned at 18-30 μm on a cryostat (Leica), and mounted on SuperFrost Plus slides (Fisher). The tissue sections were blocked for 30-60 minutes at room temperature in blocking solution (PBST with 2.5% NDS, 1% BSA, 2.5% gelatin). Sections were incubated with primary antibodies diluted in blocking solution at 4°C overnight or 1 hour at RT. Sections were then washed three times with PBS and incubated with secondary antibodies conjugated with Alexa-488, RRX or Alexa-647 (1:800 Life Technologies) in blocking solution at room temperature for 2 hours. Finally, sections were washed three times with PBS and mounted with ProLong Gold or Diamond Anti-fade Mountant (Thermo Fisher Scientific).

#### Whole-mount immunofluorescence staining

Isolated small and large intestinal tissues from mice were flushed with ice-cold PBS. Tissues were opened longitudinally and spread with the epithelial side up on Whatman filter paper or left as tubes. If left as tubes the intestinal tissue was cut under a dissection microscope (Leica) into ∼2 mm thick strips or rings containing 1-2 rows of intact crypt-villus axes to image a side view of the intestine. Mouse and human tissues were fixed with 4% PFA in PBS for 1-2 hours at 4°C, washed three times in PBS and then permeabilized for at least 3 hours in PBST followed by 1 hour in blocking solution (PBST with 2.5% NDS, 1% BSA, 2.5% gelatin). For immunolabeling, samples were incubated with primary antibodies in PBST for 24 hours at room temperature followed by 8-10 washes in PBST. Samples were incubated overnight at room temperature with secondary antibodies conjugated with Alexa-488, RRX, or Alexa-647 (1:200 Life Technologies) in blocking solution with DAPI (2 µg/ml, 1:500). Samples were washed at least 8 times with PBST containing DAPI (1:500) for 4 hours at room temperature prior to tissue clearing.

#### Tissue clearing

Ethyl-cinnamate-based tissue clearing was performed as previously described with some modifications (Gur-Cohen et al., 2019). Stained intestinal tissue was transferred through 30%, 50% and 70% ethanol diluted in UltraPure water and adjusted to pH 9.0 for 1 hour each, all on a rocker at RT. Dehydrated tissues were then incubated twice in 100% ethanol for at least 1 hour each at RT. Tissues were next transferred into 500 µl of ethyl cinnamate (Sigma) in Eppendorf tubes (Polypropylene) for clearing. Cleared intestinal tissue was mounted with ethyl cinnamate drops between 2 glass cover slips size 22×40 mm, #0 (Electron Microscopy Science), and placed in the microscope slide holder to acquire images.

#### Staining for proliferation via EdU incorporation

Staining for proliferation in tissue sections and organoids was performed according to manufacturer’s instructions using the Click-iT™ EdU Cell Proliferation Kit for Imaging, Alexa Fluor™ 647 dye (Invitrogen). To label proliferative cells in organoids, 10 µM EdU was added to ENR medium *in vitro* for 10 or 60 minutes (unless specified otherwise) before collecting organoid samples as described earlier. To label proliferative cells *in vivo*, mice were intraperitoneally injected with 123 mg/kg EdU and sacrificed after the indicated amount of time.

#### Imaging and confocal microscopy

Tissue sections, primary cells and organoid samples were imaged with the 20x objective of an AxioObserver.Z1 epifluorescence microscope equipped with a Hamamatsu ORCA-ER camera and ApoTome.2 (Carl Zeiss) slider. Additionally, organoid samples were imaged at 20x on an inverted LSM 780 laser scanning confocal microscope (Zeiss) or an inverted spinning disk confocal system (Andor Technology Inc). The LSM 780 confocal microscope is equipped with inverted Zeiss Axio Observer Z1 microscope with definite focus, 405, 440, 488, 514, 561, 594 and 633 laser lines. The system was driven by the Zen software (Zeiss). Cleared whole-mount tissue specimens were exclusively imaged using the 20x objective of an inverted Dragonfly 202 spinning disk confocal system (Andor Technology Inc) equipped with a 40 µm pinhole and a Leica DMi8 camera with AFC (20x air objective). Four laser lines (405, 488, 561 and 625 nm) were used for near simultaneous excitation of DAPI, Alexa-448, RRX and Alexa-647 fluorophores. The system was driven by the Andor Fusion software. For the majority of experiments z-stack images of 1-3 µm steps were collected. For each experiment images were acquired at identical settings.

#### Electron microscopy

Intestinal tissues were flushed with ice-cold PBS and fixed in 2% glutaraldehyde, 4% PFA, and 2 mM CaCl2 in 0.1 M sodium cacodylate buffer (pH 7.2) for >1 h at room temperature, post-fixed in 1% osmium tetroxide, and processed for Epon embedding. Ulltrathin sections (60–65 nm) were counterstained with uranyl acetate and lead citrate. Images were taken with a transmission electron microscope (Tecnai G2-12; FEI) equipped with a digital camera (AMT BioSprint29).

#### Image processing and analysis

Processing and rendering of imaging data was done in Imaris 9.5 (Bitplane) and included initial image processing, 3D-surface rendering, cell-spotting analysis as well as volumetric and distance measurements and statistical color coding of objects. 3D surface rendering and spots analysis were based on signal intensity per channel and a cell diameter estimate of 5-7 µm. Color-coding of objects was performed according to z-position, volume, area, or distance to another object as specified in the respective experiments. For the epithelial-vascular distance analyses in Figure 1 and its supplemental figures, surfaces were created for lymphatic and blood vessels as well as E-Cadherin-positive epithelial and Lgr5-positive stem cells. Surfaces of vascular structures were subsequently color-coded according to their distance from the epithelial surfaces, for which the values were calculated in Imaris by the nearest-distance-to-surface analysis. All image analyses were done at pre-defined identical settings across images and experimental replicates. Analysis of organoid area, perimeter and circularity was done in ImageJ (NIH).

#### Fluorescence-activated cell sorting (FACS) and analysis

##### Intestinal epithelial cell isolation from cultured organoids

Organoids were collected in Gentle Cell Dissociation Reagent as described above. A single cell suspension was obtained by a 20 minute incubation of organoids in TrypLE Express solution (Gibco) with DNase I (0.1 mg/ml) at 37°C with intermittent mixing every 5 minutes. Once the majority of the suspension consisted of single cells, PBS with 2% FBS was added 1:1 and the cells were centrifuged at 1400 rpm for 5 minutes. Cells were resuspended in FACS buffer (PBS with 2% BSA or FBS), filtered and counted. 10^5^ - 10^6^ cells were resuspended in FACS buffer with 1 µg/ml TruStain FcX^TM^ anti-mouse CD16/32 antibody (BioLegend) to block for 5 minutes before adding conjugated surface antibodies in FACS buffer at a 2X concentration. Cells were stained for 20-30 minutes at 4°C in the dark and washed twice in FACS buffer. Cells were resuspended in FACS buffer with DAPI (33 ng/ml) at a final concentration of 5-6 million cells/ml for subsequent fluorescence-activated cell sorting into a pre-coated tube filled with FACS buffer (for single-cell sequencing experiments) or ENR organoid medium (for *in vitro* culture of sorted cells). Cells were then spun down at 1400 rpm for 5 minutes and resuspended to reach a final concentration of 1200 cells/µl, if submitted for single-cell RNA-Sequencing, or 50-100 x 10^3^ cells/20 µl Matrigel droplet if put into culture afterwards.

#### Intestinal stem cell isolation from mouse small and large intestine

To enrich for intestinal stem cells (ISCs) in subsequent 10X-based single-cell RNA-sequencing experiments, male Lgr5-eGFP-IRES-CreERT2 mice were utilized at 8-10 weeks of age. Mice were age- and sex-matched between single-cell and spatial transcriptomic sequencing experiments. Isolation of ISCs was performed in separate experiments for the small and large intestine.

To obtain small intestinal ISCs the last ⅔ of the small intestine of each mouse were dissected and flushed 3 times with ice-cold PBS before cutting it open longitudinally. Mucus and excess villus epithelium were gently scraped off with a glass slide and the tissue was rinsed before cutting it into ∼5 mm pieces into 10 ml of ice-cold PBS in a 15 ml falcon tube. The pieces were washed several times in PBS before replacing the solution with 5 ml PBS and 10 mM EDTA pH 8.0 (Invitrogen) and incubating the tissue pieces on ice for 15 minutes with intermittent shaking every 2 minutes. The supernatant was discarded and tissue pieces were resuspended in 5 ml PBS with 5 mM EDTA pH 8.0 and incubated again on ice with intermittent shaking. After this step the supernatant was collected and filtered through a 70 µm cell strainer. The tissue was rinsed with another 10 ml PBS before being placed back into the tube. The previous incubation step was repeated 2-3 times and each supernatant was collected into separate precoated tubes as described. All fractions were spun down at 1300 rpm for 5 minutes, resuspended in 1 ml TrypLE Express (Gibco) each and incubated at 37°C for 5-10 minutes. PBS was added up to 10 ml and the cells were spun down at 1400 rpm, resuspended in FACS buffer (PBS + 2% FBS), filtered and stained for 20 minutes at 4°C in the dark. Samples were washed twice in FACS buffer and resuspended at 5-6 million cells/ml in FACS buffer with DAPI (33 ng/ml). For single-cell RNA-sequencing of the small intestine 50000 Lgr5-positive ISCs (alive, CD45-, CD31-, Lgr5-eGFP^+^ singlets) were spiked into 50000 bulk intestinal cells, which were generated from the lymphatic endothelial cell isolation of the small intestine that was done in parallel.

ISC isolation from the large intestine was performed in a similar way with modifications. The entire large intestine was taken per mouse, cleaned and processed as described. For the first incubation large intestinal tissue pieces were placed into PBS with 5 mM EDTA pH 8.0. Pieces were pipetted vigorously and allowed to settle by gravity before discarding the supernatant. Tissue pieces were resuspended in PBS with 2 mM EDTA pH 8.0 and put on a rocker at 4°C for 30 min. Pieces were shaken vigorously and the supernatant was discarded. After that serial fractions containing released crypts were collected by vigorous shaking. Fractions were spun at 1300 rpm and resuspended in TrypLE analogously to the protocol for the small intestine. For the large intestine 70 000 Lgr5-positive ISCs were spiked into 70 000 bulk intestinal cells. For all single-cell sequencing experiments cells were resuspended at a final concentration of 1200 cells/µl.

#### Intestinal lymphatic isolation from mouse small and large intestine

For bulk intestinal and lymphatic endothelial cell sorting experiments C57BL6/J mice, age- and sex-matched with mice from ISC sorting experiments, were utilized.

To isolate lymphatic endothelial cells (LECs) from the small intestine, the entire small intestine per mouse was dissected, washed and cut into ∼5 mm fragments into a 50 ml conical tube filled with PBS for additional washes. Afterwards tissue pieces were transferred into 25 ml wash solution (RPMI (Gibco) with 2% FBS, 100 µg/ml Penicillin-Streptomycin, 25 mM HEPES buffer and 2 mM L-Glutamine) with 1 mM EDTA pH 8.0 and 1 mM DTT for shaking at 225 rpm and 37°C for 15 minutes. The supernatant was discarded and tissue fragments were resuspended in 8 ml advanced DMEMF12 media with 100 µg/ml Penicillin-Streptomycin, 25 mM HEPES and 2 mM L-glutamine as well as DNase I (0.1 mg/ml), 0.25% collagenase and 2 U/ml Dispase. 2 ceramic beads were added per tube to mince the tissue during a 20 minute incubation at 37°C and 225 rpm. The solution was filtered through a 70 µm cell strainer, spun at 1300 rpm and the cell pellet was resuspended in FACS buffer. Cells were stained for 20 minutes at 4°C in the dark, washed twice and resuspended at a final concentration of 5-6 million cells/ml for sorting. For single-cell RNA-sequencing of the small intestine 25000 lymphatic endothelial cells (alive CD45-, CD31^+^, PDPN^+^, LYVE1^+^ singlets) were spiked into 50000 bulk intestinal cells from the same sample prep.

LEC isolation from the large intestine was done without the addition of ceramic beads for tissue fractionation. For single-cell RNA-sequencing of the large intestine 18000 LECs were spiked into 70000 bulk intestinal cells. For all single-cell sequencing experiments cells were resuspended at a final concentration of 1200 cells/µl.

#### Spatial transcriptomic analyses (10X Visium) of mouse small and large intestine

Spatial transcriptomics with immunofluorescence staining was performed following the guidelines by 10X Genomics. Immunofluorescence staining and imaging of intestinal tissue sections was optimized using the tissue optimization slide by 10X Genomics. For spatial gene expression analysis the distal parts of the small and large intestine from a C57BL6/J male mouse at 8 weeks of age were dissected, cleaned and cut into ∼7 mm long tubes. Tubes were directly embedded in OCT (TissueTek) in 7 mm x 7 mm cryomolds. Four tubes were embedded per block. 10 µm thick sections were taken and immediately placed onto the 10X Visium gene expression. Serial sections were used as technical replicates for small and large intestine. Tissue sections were incubated at 37°C for 1 minute followed by fixation in pre-chilled methanol at −20°C for 30 minutes. Tissue sections were blocked for 5 minutes at RT, before adding primary antibodies (1:50) and incubating at RT for 45 minutes. Sections were quickly washed 5 times before adding AlexaFluor-488, RRX- and AlexaFluor-647-conjugated secondary antibodies (1:1000) with DAPI (1:300) and incubating at RT for 30 minutes. Sections were washed 5 times and mounted in glycerol-based mounting medium. Recipes for mountant, blocking and wash buffers can be found in the 10X Visium spatial transcriptomics manual. Gene expression slides were immediately imaged on a 3DHistech Pannoramic Scanner (20x/0.8NA air objective) followed by tissue permeabilization for RNA release and cDNA synthesis. Spatial gene expression sequencing was done in collaboration with the Molecular Cytology Core and Integrated Genomics Operation at Memorial Sloan Kettering Cancer Center (MSKCC). Image processing was done using the CaseViewer Software (3DHistech). Images were manually aligned with the spot layout per capture area of the gene expression slide in Loupe Browser (10X Genomics) and exported as .json files for subsequent data processing in spaceranger. FASTQ files from sequencing (NovaSeq) were processed via spaceranger (version 1.2.1) count to align reads to the GRCm38 (mm10) reference genome and generate count matrices for subsequent bioinformatic analyses. We obtained 2,097 and 2,574 spots under tissue with median UMIs of 10,199 and 9,009 per spot for two replicates of small intestines. For the two replicates of large intestines, we obtained 2,096 and 2,180 spots under tissue with median UMIs of 19,320 and 19,814 per spot.

#### Protein secretion prediction

Genes expressed at significant levels in LECs (expressed in >20% of LECs, 2055 genes), but not expressed at a significant level in >20% of blood endothelial were analyzed to identify those likely to encode secreted proteins. Proteins with a high probability of being secreted extracellularly were identified using TargetP 1.1 (Emanuelsson et al., 2000) with a cutoff of 0.8 (80% chance of being secreted) and WoLF PSORT (Horton et al., 2007) with a cutoff score of 20 for extracellular location. This generated a list of 12 genes: *Reln*, *CCl21a*, *Sned1*, *Wnt2*, *Cp*, *Rnase4*, *Lbp*, *Mmrn1*, *Ntn1*, *Thy1*, *Rspo3*, *Il33*. For downstream analysis, we excluded *Sned1* (encodes extracellular matrix protein), *Cp* (encodes copper binding glycoprotein, involved in ion transport), *Rnase4* (encodes ribonuclease), *Lbp* (encodes lipopolysaccharide binding protein, acute phase protein), *Mmrn1* (encodes carrier protein for platelet factor V), *Thy1* (encodes a cell surface protein in immunoglobulin family) based on a low probability of being directly relevant to ISC function (Bateman et al., 2021).

#### Sequencing analyses

##### Single-cell RNA-sequencing (10X) of bulk intestinal tissue and small intestinal organoids

All single-cell RNA-seq experiments were run on a NovaSeq sequencer, and FASTQ files were generated via cellranger mkfastq (version 6.0.1.). Cellranger count was used to align reads to the GRCm38 (mm10) reference genome for intestinal organoid samples and a modified version of GRCm38 (mm10) that includes the sequence of the Lgr5-eGFP-ires-CreERT2 knockin cassette for bulk intestinal tissue samples, given that intestinal stem cells for those samples were derived from Lgr5-eGFP-ires-CreERT2 mice. Finally, cellranger count was used to generate feature-barcode matrices for subsequent bioinformatic analyses

##### scRNA-seq data analyses

*Preprocessing.* We first used CellBender (Fleming et al., 2019) to remove ambient RNA molecules from the raw count matrices with the parameter --expected-cells 5000 for the small intestine dataset, 4000 for large intestine, 800 for organoid co-cultured with LEC, and 400 for organoid alone. These parameters were chosen by inspecting the barcode rank plot generated by cellranger, following author recommendations. Defaults were used for all other parameters.

We next removed low quality cells, unexpressed genes and potential doublets from the cellbender output. Following CellBender, for the in vivo tissue dataset, we filtered cells and genes separately in each dataset. For the organoid datasets, we pooled organoid datasets from two conditions and filtered them together. For each dataset, we filtered to exclude i) genes detected in fewer than 3 cells, ii) cells with under 200 genes or 1000 UMIs, and iii) cells with a mitochondrial fraction above 15%. Mitochondrial and protein-coding ribosomal protein coding genes were also omitted from downstream analyses, as these are often a large source of spurious variance that can dominate clustering and confound deconvolution (Chu et al., 2021).

To remove doublets, we assessed the performance of both DoubletDetection (Gayoso and Shor, 2020) and Scrublet (Wolock et al., 2019) on filtered count matrices. For organoid datasets, we first split the matrices by condition. In co-cultured organoids (the organoid dataset with more cells), we observed a 57-cell cluster in which both methods detected an enrichment of doublets. DoubletDetection achieved ∼2-fold higher sensitivity (identifying 33.3% cells in this cluster as doublets, compared to 17.5% by Scrublet, with in this cluster defined as positive), at similar specificity (98.4% compared to 99.4% by Scrublet, with not in this cluster defined as negative), and we thus chose to use DoubletDetection for the organoid datasets. For *in-vivo* tissue, Scrublet yielded a more restricted set of doublets, which we chose to use due to the continuous nature of the data (DoubletDetection is known to remove transitional states at times).

We chose to use Scrublet with sim_doublet_ratio = 20 to prevent excessive removal of transitional cell types that would be critical for downstream lineage trajectory analysis. After Scrublet, 2239, 5193, 896, and 430 cells were retained for downstream analyses, with a median of 17,789, 24,316, 90,373 and 84,660 UMIs per cell in small intestine, large intestine, organoid with LEC, and organoid alone datasets, respectively. Raw counts were normalized by log_2_ (X+1) transformation, where X = library size of each cell / 10^4^.

###### Dimension reduction, clustering and annotation – organoid datasets

We performed a single round of dimension reduction and clustering on filtered cells that we pooled across organoid conditions. The pooling strategy is supported by high mixing across conditions in the KNN graph: we performed dimension reduction on log-transformed UMI counts (see preprocessing section) using PCA with 30 PCs explain 71% of variance, then computed the KNN graph at k = 10 for each cell in this PC space, and found that the majority of neighborhoods (84.1%) contain cells from both conditions.

Next, we used slalom (Buettner et al., 2017) to factor out cell cycle effects that could obscure signals from differentiation processes in downstream clustering and trajectory analyses. Specifically, cell cycle markers from Kowalczyk and Tirosh et al. (Kowalczyk et al., 2015) were converted to mouse homologs using the biomaRt package (Durinck et al., 2005, 2009). Normalized and log transformed data was used as input to slalom. G2M and S phase genes were used for initialization, employing the Gaussian noise model with pruneGenes enabled, and treating condition (LEC vs organoid alone) as a covariate. We regressed out the difference between the G2M and S phase factor scores from log-normalized gene expression, and we used residuals from this regression to perform dimension reduction, clustering and trajectory analysis.

For downstream analysis, we first selected the top 5000 highly variable genes in each dataset using the scanpy function pp.highly_variable_genes, with raw counts as input, n_top_genes = 5000, flavor = ‘seurat_v3’ and span = 1. For these genes, we performed dimension reduction over the residuals of the cell cycle regression, using the scanpy.tl.pca function with the arpack solver. We selected 30 PCs, which explain 71% of variance. For clustering, we used Phenograph (Levine et al., 2015) with k = 30 and clustering_algo = ‘leiden’, which generated 11 clusters in total.

We used known cell type markers to annotate the clusters. Five clusters expressed strong enterocyte markers (e.g. *Alpi*, *Ephx*, *Apoa1*), 4 expressed stem cell and transient amplifying cell (TA) markers, (e.g. *Lgr5*, *Olfm4*, *Axin2*, *Top2a*, *Tubb5*) and the remaining 2 expressed goblet/Paneth (e.g. *Lyz*, *Muc2*) and enteroendocrine (e.g. *Chga*, *Chgb*) markers respectively (Figure S3H). To assign enterocyte clusters, we isolated these cells for analysis, and computed the z scores of markers of each enterocyte zone reported in Moor et al. (Moor et al., 2018) then averaged z scores across markers of each zone (Figure S3H). Two clusters exhibited high scores for zone 2 marker genes and were labeled bottom-like 1 and bottom-like 2; two clusters with highest zone 3 marker scores were labeled mid-like 1 and mid-like 2; and the cluster with highest zone 5 marker scores was labeled top-like.

###### Dimension reduction, clustering and cell type annotation – in vivo datasets

We performed an initial round of dimension reduction and clustering in the *in vivo* tissue data, and then zoomed in on heterogeneous clusters for additional rounds of refinement. For initial clustering, we selected the top 5000 highly variable genes in each dataset using the scanpy function pp.highly_variable_genes with raw counts as input, n_top_genes = 5000, flavor = ‘seurat_v3’ and span = 1. We applied PCA to these genes using the scanpy.tl.pca function with the arpack solver. We selected 30 PCs, which explain 63.6% and 63.1% of variance in small and large intestine datasets, respectively, and were able to capture coarse phenotyping of main cellular compartments. For clustering, we used Phenograph with k = 15 and clustering_algo = ‘leiden’, which generated 27 and 35 clusters in the small and large intestine datasets, respectively.

The *in vivo* data cover a broad spectrum of immune, stromal, endothelial and epithelial lineages. Thus, cell types of similar lineage may not cluster accurately with 30 PCs, prompting us to perform a second round of clustering among cell types from similar lineages. During this refinement stage, we selected the top 3000 highly variable genes for each subset of cells (described below), and performed PCA with 30 PCs. We then re-clustered cells using Phenograph with k = 30. The higher k value in this round was chosen to prevent over-clustering. Over-clustering can cause the reference cell states used for deconvolution (in spatial transcriptomics analysis) to be highly linearly dependent, resulting in a possible identifiability issue. To further reduce the linear dependency between clusters, we also regrouped subclusters that express similar marker genes profiles (see below).

We re-clustered cells for each dataset as follows. For small intestine, we pooled and re-clustered the stem, TA and mature enterocyte clusters (N = 848), then further divided and separately re-clustered mature enterocytes (N = 428) and stem/TA cells (N = 420). For each of the 9 resulting mature enterocyte clusters, we compared mean z scores between bottom and top zone landmark genes (Moor et al., 2018) (Figure S4E). Six clusters with lowest top-to-bottom ratio were labeled bottom zone enterocytes (N = 325); two with marginally higher ratios were labeled mid zone enterocytes (N = 63); and one cluster with substantially higher ratio was labeled top zone enterocytes (N = 40). The stem/TA group (N = 420) was refined into 7 clusters; three lacking Lgr5 expression were labeled TA (N = 191); two with high expression of Lgr5 and other ISC markers was labelled Lgr5+ stem; (N = 121); and two with highest TA marker expression, including *Stmn1* and *Tubb5*, and moderate *Lgr5* were labeled Lgr5+ progenitor (N = 108). The goblet cell group (N = 216) was refined into 5 clusters; one expressing Paneth, goblet and secretory progenitor cell markers was labeled secretory progenitor (N = 35); two with high *Mki67* expression was labeled goblet_cycling (N = 79); and the remaining two clusters were labeled goblet_1 (N = 46) and goblet_2 (N = 56). Refinement of cells expressing stromal marker genes (N = 84) yielded three sub-clusters; one with high myofibroblast marker expression was labeled as such (N = 14); and two that do not perfectly match known cell types were labeled str1 (N = 38) and str2 (N = 32), which may represent a mixture of stromal cell types.

For the large intestine dataset, we first reclustered a population expressing high stem and enterocyte and low goblet cell markers (N = 2187), yielding 16 clusters. Four clusters exhibit high enterocyte and low stem and TA marker expression; two of these have high crypt top enterocyte marker expression and were labeled mature enterocyte 1 (N = 49) and 2 (N = 102); and the others two were labeled enterocyte 1 (N = 72) and 2 (N = 78). Eleven clusters exhibit high expression of *Lgr5* and other stem cell markers such as *Ascl2* and *Smoc2*; six of these have high cell cycle marker expression and were labeled Lgr5+ stem cycling (N = 815); and the remaining five were labeled Lgr5+ stem (N = 946). The final cluster has higher TA but intermediate stem marker expression, and was labeled TA (N = 125). Reclustering of cells expressing goblet markers (N = 1577) yielded 10 clusters; four similar clusters that express high cell cycle markers were labeled goblet cycling (N = 486); we combined one small cluster goblet_2 (N = 16) with goblet_8, the cluster with highest Pearson correlation, generating the group goblet_2_8 (N = 246); and the remaining four clusters were labeled by Phenograph cluster ID. Reclustering of stromal cells (N = 473), generated 8 clusters; three with high glial cell, pericyte and myofibroblast marker expression were labeled with the corresponding cell type (N = 12, 17 and 91, respectively); three clusters express str1, 2, and 3 marker genes from Kinchen et al. (Kinchen et al., 2018) and were labeled accordingly (N = 116, 65 and 172, respectively); and we excluded a small cluster (N = 30) with low library size and number of detected genes, which likely reflects low quality cells, from trajectory and deconvolution analyses.

###### Marker genes used in Figure S3 and Figure S4 for cell type identification

To identify cell types in the organoid sequencing data, gene sets highlighting differentiated enterocyte subsets along the intestinal villus axis (Moor et al., 2018) and those specifying remaining epithelial populations, i.e. stem/TA, goblet/Paneth and enteroendocrine cells (Biton et al., 2018), were taken from the respective literature. Cell type annotation in the *in vivo* sequencing data was done based on established marker gene sets to identify cells of the epithelial (Haber et al., 2017), stromal (Kinchen et al., 2018) and immune (Xu et al., 2019) composition present in the mouse intestine.

###### Differential abundance analysis – organoid dataset

We adapted Milo (Dann et al., 2021) to test for the differential abundance of cells within defined neighborhoods, between two conditions lacking replicates. The preprocessed, cell-cycle-corrected organoid datasets (above) were used as input by Milo. We first used the buildGraph function to construct a KNN graph with k = 30, using 30 principle components (d = 30). Next, we used the makeNhoods function from Milo function to assign cells into neighborhoods based on their connectivity over the KNN graph. For computational efficiency, we subsampled 10% of representative cells, i.e. the index cells, from the KNN graph to define neighborhoods using Milo’s min-max sampling scheme, originally proposed in (Setty et al., 2016). This step generated 95 neighborhoods. Next we tested for differences in the abundance of cells from each condition in each neighborhood. As our data lacks replicates, we modified the negative binomial test used by the original Milo implementation to a two-sided Fisher’s exact test. Lastly we corrected for multiple testing for p values outputted by Fisher’s exact tests to account for the overlap between neighbourhoods using the Milo function graphSpatialFDR. Specifically, the p value of each neighborhood was corrected using the Benjamini–Hochberg method, but weighted by the reciprocal of the neighborhood connectivity. The connectivity was measured as the Euclidean distance calculated in the PC space (d = 30) to the *k*th neighbor of the index cell that defines each neighborhood with k=10. Additionally we computed the log_2_ fold change of number of cells between the LEC co-\culture vs. the organoid alone condition. The spatial FDR and log_2_ fold change of number of cells between two conditions in each neighborhood was used for visualization.

###### Differential gene expression analyses for the organoid dataset using Mast

We performed differential expression analysis using Mast (Finak et al., 2015) on organoid data using the preprocessed log normalized expression data described above. We first subsetted the cells from the enterocyte clusters in each condition, and fitted the zero-inflated regression model using the zlm function of the Mast R package. We performed a likelihood ratio test over two linear models to derive the log_2_ fold change and statistical significance of change between the gene expression from two culture conditions, i.e. LEC coculture vs. organoid alone. The alternative model used two covariates. The first covariate is a categorical variable which represents the culture condition. The second covariate is a continuous variable representing the cellular detection rate which was calculated as the number of cells with zero expression followed by scaling to mean of zero and standard deviation of one. Only the second covariate is included by the null model. Multiple testing correction was performed using false discovery rate. We defined genes significantly changed between two conditions as those with FDR less than 0.01, and log_2_ fold change greater than 0.1. The log_2_ fold change of genes significantly differentially expressed were visualized for each zone marker genes in Figure 2F.

##### Trajectory analysis for organoid data

To understand the emergence of the different cell population frequencies between organoids grown alone or in co-culture with LECs, we used features of the CellRank package (Lange et al., 2020) to map potential differentiation trajectories separately for each culturing condition. We chose to supply CellRank with the CytoTRACE kernel (Gulati et al., 2020) due to its superior performance quantifying pseudotemporal ordering specifically in stem and very early progenitor populations. We used cell cycle-corrected gene expression (described above) as input to our trajectory analysis. First, we constructed the pseudotemporal ordering of cells using CytoTRACE over genes with greater than 2 reads. We found that applying such filtering was important, as it removed genes that were not confidently expressed but still detected at a high sequencing depth in our organoid data. Second, we constructed the undirected KNN graph using the Palantir (Setty et al., 2019) kernel, since it is particularly effective at modeling both rare and frequent subpopulations, as well as handling trajectories at different scales. Third, transition probabilities between cells in the opposite direction of the CytoTRACE ordering were pruned away from the transition matrix by setting the parameter threshold_scheme of the compute_transition_matrix function to “hard”. Lastly, a transition matrix was constructed based on the ordered KNN graph and projected onto the force directed layout for visualization. To predict the distribution of terminal states, we simulated 100 random walks from the transition matrix in the forward direction starting from cells randomly sampled from the stem/TA population in each condition. We used the default parameters of the plot_random_walks function, which simulated the number of random walk steps equal to 25% of the total number of cells in each condition.

#### BayesPrism algorithm

BayesPrism is a fully Bayesian deconvolution method that jointly models the cell type fractions and cell type-specific gene expression from each mixture using scRNA-seq as the reference. It was previously developed for deconvolving bulk RNA-seq, and we show here that it represents a general deconvolution framework that naturally extends to spatial transcriptomic data. For spatial transcriptomic data, BayesPrism takes 3 inputs, the raw count matrix of scRNA-seq, cell type labels of each cell and the raw count of each spot from the Visium data. The first two inputs were derived from the preprocessing and clustering/cell type annotation steps of the *in vivo* scRNA-seq data described above. The high-level details of the algorithm are as follows. The scRNA-seq count matrix was collapsed to generate the gene expression profile matrix φ, by summing over the cells in each cell type and then renormalizing such that the sum across genes is 1. The matrix φ which essentially describes the multinomial parameters for the count matrix of each cell type. It is used as the prior information to infer the posterior distribution of cell type fractions and gene expression conditional on the observed expression in each spatial spot:

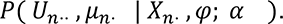

*U*_*nsg*_ represents the number of counts assigned to the *g_th_* gene of *s_th_* cell type in the *n_th_* spot. *μ*_*ns*_ represents the estimates of fraction of counts from the *s_th_* cell type and the *n_th_* spot, an estimate assumed to be proportional to the fraction of cells in each spot. *X_ng_* represents the counts of the *g_th_* gene in the *n_th_* spot, measured by spatial transcriptomics. *α* is a non-informative weak dirichlet prior hyper-parameter set to a small value=10^-8^ by default. BayesPrism implements an efficient Gibbs sampling scheme to sample *U* and *μ* from the posterior distribution, followed by marginalizing each other and then summarizing using the posterior mean. We also leveraged the information shared across thousands of spatial spots in each dataset to update *φ* using *U*⋅, to construct the updated gene expression profile *ψ* of the same dimension as *φ*, by setting the argument first.gibbs.only=FALSE in the run.Ted function provided by BayesPrsim. This procedure allows for better accommodating biological variation between the reference scRNA-seq data and the spatial transcriptomics. *ψ* was then used to re-estimate the cell type fractions using the posterior distribution described above, while marginalizing *U*:

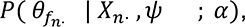

where *θ_fns_* is the updated version of *μ_ns_*, which also represents the estimates of fraction of counts from the *s_th_* cell type and the *n_th_* spot. *θ_f_* was used for all downstream analysis.

BayesPrism has been mathematically shown to be intrinsically invariant to linear batch effects while being robust to other types of non-linear batch effects and biological variations. As all other methods developed for deconvolving spatial transcriptomic data require a large number of spatial spots to learn the linear correction factor for batch effects between scRNA-seq and spatial transcriptomics, we show for the first time that directly modeling such linear batch effects is not necessary (see benchmarking described below). More importantly, while most spatial deconvolution methods only supply cell-type fractions, BayesPrism supplies both cell type fractions and cell type specific gene expression for each spatial transcriptomic spot. Therefore, BayesPrism represents a powerful deconvolution tool for spatial transcriptomics.

#### Cell type deconvolution using BayesPrism

We deconvolved the Visium data of small and large intestine dataset using the corresponding *in vivo* scRNA-seq data. To increase signal to noise ratio for deconvolution, we selected marker genes that are differentially upregulated in each cell type. We then took the union of these marker genes and deconvolved over these genes. Specifically, we performed the pairwise t-test using the findMarker function from the scran package (Lun et al., 2016) between each pair of cell types over the log transformed reads output by our scRNA-seq pipeline. We then required both the maximum p value to be less than 0.05, and the minimum log_2_ fold change to be above 0.05. This generated a total of 3,265 genes for the small intestine dataset and 3,588 genes for the large intestine dataset. To also deconvolve the gene expression profiles for known receptor and ligand genes, we also added 1,349 receptor ligand genes compiled from multiple sources (Baccin et al., 2019; Jin et al., 2021), totalling up to 4,266 and 4,487 genes for the small and large intestine dataset. Additionally we excluded spots with low UMI counts from deconvolution. Spots with less than 2000 UMIs in the small intestine dataset and 4000 UMIs in the large intestine dataset were removed. Each biological replicate was deconvolved separately. To ensure a fair comparison between deconvolution methods, we used the same set of genes and the same set of spatial spots in benchmarking.

#### Benchmarking Deconvolution of Spatial Transcriptomics

A big challenge in the deconvolution field is the lack of ground truth for benchmarking. To estimate some ground truth of the cell type fractions from the Visium data we used a simple marker genes based approach. We defined markers for each cell type based on a stringent set of differentially expressed genes, using scran’s findMarker function (Lun et al., 2016), but with a more stringent criterion. Specifically we used the top 50 differentially expressed genes ranked by log_2_ fold change compared to other cell types. Additionally, only genes with maximum p value less than 0.01, and the minimum log_2_ fold change above 0.1 were used as markers for estimating cell type fractions. To estimate the fraction of each cell type in each spatial spot, we calculated the fraction of reads expressed in each gene set over all marker genes selected. While this estimate is not a ground truth, we took it as a reasonable estimate and used it for comparison in Figure S5A.

For our benchmarking comparison, we executed other methods as follows: For Stereoscope, we trained the scRNA-seq data with 1000 epochs, and the deconvolution model with 10000 epochs until convergence. For cell2location, we trained the scRNA-seq data by RegressionModel using batch_size=2500, train_size=1, lr=0.002, max_epochs=40000 (small intestine), and 30000 (large intestine), followed by estimating the cluster mu by simulating 1000 samples from the posterior distribution using the same batch size. The deconvolution model was trained with the hyper-parameters N_cells_per_location=30, detection_alpha=200, max_epochs=30000, batch_size=None and train_size=1. The same setup as the scRNA-seq model was used to draw samples from the posterior distribution to estimate the cell abundance score. We used the 5% quantile of this posterior distribution, representing the cell abundance scores with high confidence. Lastly we normalized the abundance score to sum to one for each spatial spot. For destVI, we trained the scRNA-seq model using max_epochs=250, and the deconvolution model using max_epochs=2500 until convergence. For RCTD, we used the parameter doublet_mode = ‘full’ to output the fraction of every cell type without forcing a constraint to make a fair comparison with other tools. For BayesPrism, all parameters were at their default values.

#### SpaceFold cartography

*Projecting spots onto a 1D axis:* The Visium data only comprised ∼4 spots for each crypt. To achieve higher spatial resolution we used the following features of our data: (1) Our spatial transcriptomics data consisted of hundreds of crypts. (2) Each crypt is organized in a highly stereotypical fashion. (3) The spatial spots along each crypt are each shifted in a slightly different position (see Fig 5A, left). Given the stereotypical nature of the crypt, we assume that two spots with similar cell type fractions are in a similar spatial position along the crypt, and moreover these cell type fractions change gradually along the crypt axis. Therefore, we describe each spatial transcriptomic spot using a vector representing the cell type abundances as computed by BayesPrism (in this application ∼30 cell types). In the next step, we aim to use non-linear dimensionality reduction to reduce the 30-dimensional cell-type fraction vector into a 1D axis representing position along the crypt axs. We used the cell type fraction inferred by BayesPrism as the input, and used PHATE (Moon et al., 2019) to project this vector onto a 1D axis. We chose PHATE due to its ability to model lighter tails in the kernel function using the α-decaying kernel, which decays faster than the common Gaussian kernel. A faster decaying kernel can be beneficial in creating a step function-like window with high transition probabilities only for spots with similar cell type compositions. Such narrow windows prevent spots from regions far away from each other, such as those from the crypt base and crypt top, from having high transition probabilities to induce false edges in the transition matrix.

With thousands of spots, we were able to project them onto the 1D crypt axis continuously at a much higher spatial resolution than the original Visium spots. Moreover, with BayesPrism’s cell-type specific gene expression, provided for each spot, we can plot cell-type specific gene trends along this axis (see section below for details). While we chose to use BayesPrism for computation of cell type fractions in each spot and PHATE for non-linear dimensionality reduction, alternative spatial deconvolution and dimensionality algorithms can be used within the SpaceFold framework.

To run SpaceFold on our data, we first compute the z-score of the fraction of each cell type by centering to mean zero followed by scaling to standard deviation. This ensures that the distance calculated by the transition matrix will not be dominated by cell types of high abundance. To run PHATE, we set knn=10 and ndim=1, and other parameters at their default values, which includes a high α value, α=40. The PHATE embedding was rescaled to between 0 and 1 for visualization. Intestine tissue is known to show heterogeneous gene expression across disparate tissue segments (e.g. proximal or distal) (Parigi et al., 2021) which may violate the assumption of the identical and repetitive unit. This may confound PHATE, causing the one dimensional axis unable to fully model the underlying structure. Indeed, in the visium samples of large intestine tissue, two of the three tissue regions displayed differences in cell type abundance compared to the third. To ensure the model assumption of SpaceFold holds, we run SpaceFold separately on spots from each region. We found the predicted 1D axis correlated well between two regions, and hence merged them for visualization in Figure 4C.

Although SpaceFold cartography was developed for mapping gene expression along the crypt axis, it can be easily generalized to other tissue types of repetitive structures. For tissues of more complex structures, projecting onto a 2D plane is also an option.

##### Visualizing gene expression trends along the projected axis

BayesPrism outputs a deconvolved 3-dimensional gene expression tensor *U_nsg_* (see the section BayesPrism algorithm) with the number of reads of each gene *g* in each cell type *s* from each spot *n*. To compare gene expression in one particular cell type between different spots, deconvolved reads need to be normalized with respect to the total number of reads in each cell type. One naive way of normalization would be to first compute a normalizing factor by summing over the dimension *g* over *U_nsg_* to derive a size factor *U_ns_* which represents the total number of reads contributed by each cell type to each spot, followed by dividing *U_nsg_* by the normalizing factor to obtain the normalized expression <u>*U*</u>*_nsg_*:

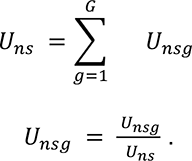

One issue with the such normalization is *U_ns_* will approach zero for spots that contain low or no cell type *s*, and hence cause the normalization to be unstable. To circumvent this, we compute normalized gene expression only for spots showing a substantial amount of the cell type of interest. We used the mixture model with 2 components to separate spots that contain cell types of interest from the background. Specifically we used the Gaussian mixture model to fit the reads contributed by each cell type to each spot, *U_ns_*, using the mclust R package (Scrucca et al., 2016), and used the Gamma mixture model to fit cell type fraction, *θ_fns_*, using the mixtools R package (Benaglia et al.) for each cell type *s*. Spots from the clusters of higher mean from the Gaussian mixture and the Gamma mixture models were intersected to represent spots containing the cell type of interest. Due to the sparsity of Visium data, we grouped the spots selected for each cell type into 20 bins according to their projected 1D axis, and take the mean and standard error of the normalized gene expression values *U_nsg_* over each bin. For visualization, we fitted a local 2nd order polynomial regression curve for the mean, and mean ± 2 standard error against the mean of 1D coordinates of each bin using the R package loess (Cleveland et al., 2017) with the smoothing parameter “span=0.75”.

To visualize the gene expression for zone markers we plot the mean z-score of normalized gene expression in enterocytes over the marker genes as follows (Figure 5E). For each zone marker genes *g* selected for deconvolution, we first summed *U_nsg_* over *s*, for *s* belong to bottom zone, mid zone and top zone enterocytes, and divide the sum by the total UMI of each spot to get a normalized expression for gene *g* in spot *n* in all enterocytes. We then computed the z score for each gene, and took the mean of the z scores over the markers within the list of each zone. Taking the mean of z-score makes the expression comparable between different zones. We grouped all spots into 50 bins according to their projected 1D axis, and took the mean and standard error of the mean z-scores computed above over each bin. The local 2^nd^ order polynomial regression curve was computed in the same way as described above.

To quantify the total gene expression level from a group of cell types of different spatial distributions, such as all Lgr5^-^ cells, as shown in left and center panels in Figure 5D, we report the unnormalized sum of *U_nsg_* over the cell types from the selected group. We did not normalize by *U_ns_* as we found that the difference in the spatial distribution of cell types sometimes may inflate denominators. This will be the case when, for example, a cell type that does not express the gene of interest is present at a high abundance over a region of spots.

#### Statistics and reproducibility

Statistical analyses were performed using the GraphPad Prism 8 software. All *in vitro* experiments were repeated at least 3 times in duplicates and representative data are shown. Technical replicates were also used for spatial gene expression analysis. For *ex vivo* tissue stainings tissue from at least two age- and sex-matched mice was utilized per experiment. The number of mice utilized for imaging studies is indicated as *n* = x mice per timepoint or condition unless indicated otherwise. Phenotypic analyses of organoids were performed independently by 2 experimenters in a blinded manner. Unpaired two-tailed Student’s *t*-tests were used to ascertain statistical significance between two groups. One-way analysis of variance (ANOVA) was used to assess statistical significance between three or more groups with one experimental parameter. See figure legends for more information on statistical tests.

## Notes

### Competing Interest Statement

The authors have declared no competing interest.

